# StrVCTVRE: A supervised learning method to predict the pathogenicity of human genome structural variants

**DOI:** 10.1101/2020.05.15.097048

**Authors:** Andrew G. Sharo, Zhiqiang Hu, Shamil R. Sunyaev, Steven E. Brenner

**Author notes:** Email: Andrew G. Sharo, Zhiqiang Hu, Shamil R. Sunyaev, Steven E. Brenner.

## Abstract

**Background:** Whole genome sequencing resolves many clinical cases where standard diagnostic methods have failed. However, at least half of these cases remain unresolved after whole genome sequencing. Structural variants (SVs; genomic variants larger than 50 base pairs) of uncertain significance are the genetic cause of a portion of these unresolved cases. As sequencing methods using long or linked reads become more accessible and structural variant detection algorithms improve, clinicians and researchers are gaining access to thousands of reliable SVs of unknown disease relevance. Methods to predict the pathogenicity of these SVs are required to realize the full diagnostic potential of long-read sequencing.

**Results:** To address this emerging need, we developed StrVCTVRE to distinguish pathogenic SVs from benign SVs that overlap exons. In a random forest classifier, we integrated features that capture gene importance, coding region, conservation, expression, and exon structure. We found that features such as expression and conservation are important but are absent from SV classification guidelines. We leveraged multiple resources to construct a size-matched training set of rare, putatively benign and pathogenic SVs. StrVCTVRE performs accurately across a wide SV size range on independent test sets, which will allow clinicians and researchers to eliminate about half of SVs from consideration while retaining a 90% sensitivity.

**Conclusions:** We anticipate clinicians and researchers will use StrVCTVRE to prioritize SVs in patients where no SV is immediately compelling, empowering deeper investigation into novel SVs to resolve cases and understand new mechanisms of disease. StrVCTVRE runs rapidly and is available at https://compbio.berkeley.edu/proj/strvctvre/.

## Background

Whole genome sequencing (WGS) can identify causative variants in clinical cases that elude other diagnostic methods(1). As the price of WGS falls and it is used more frequently, researchers and clinicians will increasingly observe structural variants (SVs) of unknown significance. SVs are a heterogeneous class of genomic variants that include copy number variants such as duplications and deletions, rearrangements such as inversions, and mobile element insertions. While a typical short-read WGS study finds 5,000–10,000 SVs per human genome, long-read WGS is able to identify more than 20,000 with much greater reliability(2–4). This is two orders of magnitude fewer than the ∼3 million single nucleotide variants (SNVs) identified in a typical WGS study. Still, despite their relatively small number, SVs play a disproportionately large role in genetic disease and are of great interest to clinical geneticists and researchers(5, 6).

SVs are of clinical interest because they cause many rare diseases. Most SVs identified by WGS are benign, but on average, a given SV is more damaging than an SNV due to its greater size and ability to disrupt multiple exons, create gene fusions, and change gene dosage. In a study of 119 probands who received a molecular diagnosis from short-read WGS, 13% of cases were caused be an SV(7). Similarly, an earlier study that found 7% of congenital scoliosis cases are caused by compound heterozygotes comprised of at least one deletion(8). Yet, since SVs continue to be challenging to identify and analyze, these figures may underestimate the true causal role that SVs play in rare disease. Indeed, in some rare diseases, the majority of cases are caused by SVs. For example, deletions cause most known cases of Smith-Magenis syndrome, and duplications cause most known cases of Charcot-Marie-Tooth disease type 1A(9). This suggests that for rare disorders, SVs constitute a minor yet appreciable fraction of pathogenic variants.

To continue discovering SVs which cause disease, researchers face a daunting challenge: prioritizing and analyzing the tens of thousands of SVs found by WGS. Best practices for SV prioritization are evolving, and generally mirror steps used to prioritize SNVs. Few SV-tailored impact predictors have been developed, but a small number of published studies have focused on identifying pathogenic SVs from WES(10, 11) and WGS(7, 12, 13) and have identified a handful of important steps. Removing low-quality SV calls is essential, as short-read SV callers rarely achieve precision above 80% for deletions and 50% for duplications, even at low recall(14). Most studies remove SVs seen at high frequency in population databases or internal controls(6, 15). Moreover, many studies only investigate SVs that overlap an exonic region, as non-coding SVs remain particularly difficult to interpret. Depending on its sensitivity, a pathogenic SV discovery pipeline may produce tens to hundreds of rare exon-altering SVs per proband to be investigated. These values are consistent with a recent population-level study that estimates SVs comprise at least 25% of all rare predicted loss-of-function events per genome(16). Prioritizing SVs will be necessary for the majority of probands, as shown by a study of nearly 500 unresolved cases that found one or more SVs that warranted further investigation in 60% of cases(7). Clinically validating all SVs of uncertain significance in a genome is currently infeasible, and cohort size for rare diseases will likely never reach a scale sufficient to statistically associate these SVs with disease. Therefore, computational tools are needed to prioritize and predict the pathogenicity of rare SVs.

Among methods that consider SVs, several annotate the features of SVs but very few prioritize SVs by pathogenicity. General-purpose annotation frameworks such as Ensembl’s Variant Effect Predictor (VEP)(17) and SnpEff(18) both annotate SVs with broad consequences based on sequence ontology terms (e.g., transcript_ablation), which we found are not sufficient for effective prioritization. One standalone annotator, SURVIVOR_ant(19), annotates SVs with genes, repetitive regions, SVs from population databases, and user defined features. This and similar tools put the onus on researchers to provide informative features and determine how to consider these features in combination, a difficult challenge. A complementary approach is to annotate SVs using cataloged SVs known to be pathogenic or benign. One such SV annotator, AnnotSV(20), ranks SVs into five classes based on their overlap with known pathogenic or benign SVs and genes known to be associated with disease or predicted to be intolerant to variation. This approach can be successful when a disease-causing SV has previously been seen in another proband and was cataloged as pathogenic, but we show it has limitations when a disease-causing SV is novel. In contrast, SNVs can be effectively prioritized by methods such as Revel(21) and VEST(22) that integrate diverse annotations to provide a quantitative score. Similarly powerful methods are needed to predict SV pathogenicity.

In order to provide a summary pathogenicity score to prioritize rare SVs genome-wide, a predictor must address two questions. The first question is whether a gene is likely associated with a Mendelian phenotype. This relationship can be predicted through gene importance features. The second question is whether an SV impacts gene function, which requires considering intragenic features. Although these are two separate questions, for convenience researchers often combine them into a single summary score. Few methods provide such a summary score for SV pathogenicity. One standalone impact predictor, SVScore(23), calculates the deleteriousness of all possible SNVs within each SV (using CADD(24) scores by default), while considering SV type and gene truncation. SVScore then generates a summary score by aggregating across these CADD scores (mean of the top 10% by default), and this approach has shown promise in identifying SVs under purifying selection(23). Another stand-alone predictor, SVFX(25), integrates multiple features into a summary score, but focuses on somatic SVs in cancer and germline SVs in common diseases so we do not discuss it further.

In this manuscript, we introduce StrVCTVRE (<u>Str</u>uctural <u>V</u>ariant <u>C</u>lassifier <u>T</u>rained on <u>V</u>ariants <u>R</u>are and <u>E</u>xonic), a method that generates a summary pathogenicity score for exon-altering SVs. We anticipate clinicians and researchers will use StrVCTVRE to prioritize rare SVs associated with Mendelian phenotypes. Since nearly all pathogenic SVs are rare (minor allele frequency (MAF) < 1%), the salient challenge in resolving undiagnosed cases is to distinguish rare pathogenic SVs from rare benign SVs(16). Existing SV predictors have been trained and assessed on common benign SVs(23, 25), so they may rely on features that instead separate common SVs from rare SVs and may not be optimal for this clinical question(26). StrVCTVRE is the first method trained to distinguish benign rare SVs from pathogenic rare SVs. StrVCTVRE is available at https://compbio.berkeley.edu/proj/strvctvre.

## Results

### StrVCTVRE design and assessment

StrVCTVRE is implemented as a random forest, in which many decision trees ‘vote’ for whether a given SV is pathogenic. The StrVCTVRE score reflects the fraction of decision trees that ‘voted’ that the SV is pathogenic. The decision trees are shaped by a learning algorithm, in which each tree sees thousands of example SVs from a training dataset of known pathogenic and benign SVs, and the decision nodes are optimized for accuracy. To promote diverse trees, each node of the decision tree uses only a random subset of the features. Finally, StrVCTVRE is assessed on a held-out test dataset and independent test datasets.

### Characterization of StrVCTVRE features

To classify SVs, StrVCTVRE employs 17 features in five categories: gene importance, conservation, coding sequence, expression, and exon structure of the disrupted region (see Methods, Table 2 for details). We assessed gene importance using two features that summarize the degree of depletion of predicted loss-of-function (pLoF) variants in healthy individuals: pLI(27) and LOEUF(15). Although LOEUF is effectively an updated, continuous version of pLI, and the two are highly correlated, we found better performance when both were included rather than just one. To explicitly capture when an important gene is highly impacted by an SV, we included two additional features: pLI of a highly impacted gene and LOEUF of a highly impacted gene. We define a gene as highly impacted when an SV overlaps the APPRIS(28) principal start codon or 50% of CDS. To specifically model coding sequence (CDS) disruptions, we used three coding features: percentage of the CDS overlapped by the SV, distance from the CDS start to the nearest position in the SV, and distance from the CDS end to the nearest position in the SV. We included a single conservation feature, phyloP of 100 vertebrates(29), by considering the average of the 400 most conserved sites in the SV. PhyloP produced the best classification among the conservation features we investigated (see Methods) and was the most informative conservation feature in a rare missense variant classifier(21). To infer expression impacts from the SV, we included the average expression across all tissues for each exon in the SV, the proportion of gene transcripts that included each exon in the SV, and the overlap with known topologically associating domain (TAD) boundaries. To model potential differences that drive the pathogenicity of deletions and duplications, we included as a feature whether an SV is a deletion or duplication. The remaining features were related to the structure of exons in the SV including the number of exons in a disrupted gene, the number of exons disrupted, whether any affected exons were constitutive, whether all disrupted exons could be skipped in frame, and the order of the exon in the transcript. When multiple exons or genes were disrupted, we typically took the value of the most severely impacted one, as appropriate (see Methods). Missing or non-applicable feature data were replaced by the median value of each feature.

### Correlation and relative importance of SV features in StrVCTVRE

Clusters emerged when we calculated these features for our SV training set, computed the correlation between each feature, and clustered by correlation (Fig. 1a). The most prominent cluster (labeled i) contains gene importance, conservation, CDS, and one exonic feature, with most correlations above Spearman’s ρ = 0.6. Since both gene importance of highly impacted gene features are present in this cluster, the other features in this cluster may also capture when an important gene is highly disrupted. A smaller cluster (labeled ii) included the remaining gene importance features, pLI and LOEUF. Expression features and deletion/duplication status were the features least correlated with all other features (all ρ ≤ 0.26). This low correlation suggests that these features capture unique information, which is unsurprising for deletion/duplication status. But given the relative importance of some expression features (Fig. 1b), our results suggest expression data contains both orthogonal and valuable information for determining SV pathogenicity. The two features capturing gene importance of a highly impacted gene were the features most correlated with each other (ρ = 0.97), indicating that pLI and LOEUF are generally interchangeable for assessing the importance of highly disrupted genes.

**Fig. 1.**
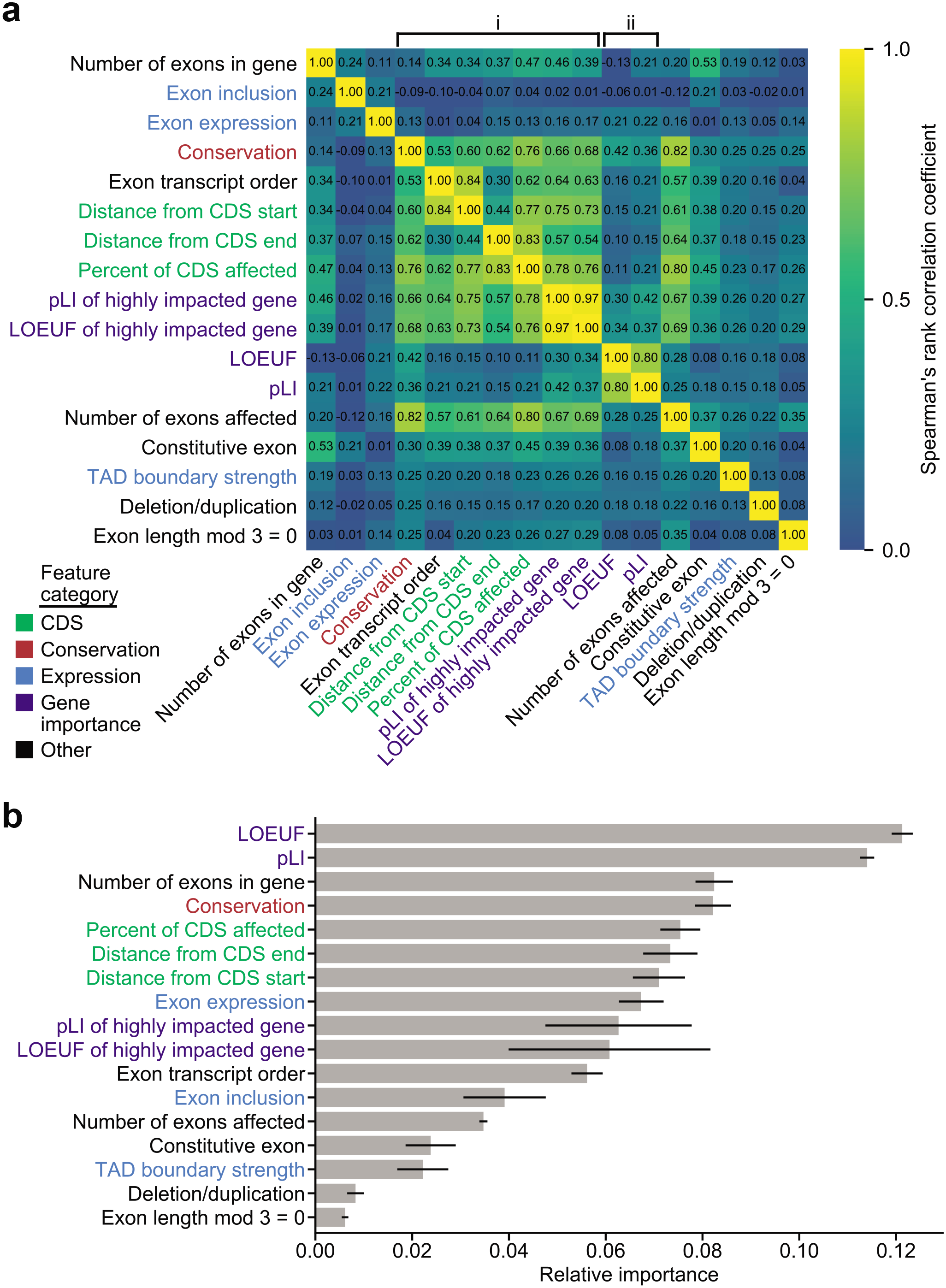
By considering feature clustering and importance, we can identify features providing unique and predictive information. **a** Correlation matrix of StrVCTVRE features in training data. Features were ordered by hierarchical clustering, and some values were reversed to reduce negative correlation between features. Values represent Spearman’s rank correlation between features. Text is colored by feature category. **b** Feature importance of StrVCTVRE features. Gray bars indicate feature importance, estimated using mean decrease in impurity (Gini importance). Black lines indicate 95% confidence intervals. Note that exon expression had high importance yet was uncorrelated with all other features, suggesting it captures unique and predictive information.

By training on thousands of example SVs, StrVCTVRE discovers which features are useful for discriminating between pathogenic and benign SVs (Fig. 1b). Using Gini importance (see Methods), we found gene importance features were most useful to StrVCTVRE. This was followed by a group of features with similar importance that include the number of exons in a gene, conservation, CDS features, exon expression, and gene importance of a highly impacted gene. The value of these features is largely intuitive; gene importance, CDS, and conservation features are expected to be helpful to assess pathogenicity. In contrast, we suspect number of exons in gene is highly ranked due to sampling bias. We found that many well-studied pathogenic genes have numerous exons (DMD, NF1, BRCA2), and these genes have many representative SVs in our dataset even after removing near-duplicates (Fig. S1, Methods). This may lead StrVCTVRE to have improved performance on these known clinically relevant genes, but reduced performance genome-wide (discussed further below). Surprisingly, several exonic features had relatively low importance, which may have been caused by the sparsity of SVs in our dataset that alter just a single exon. The low importance of TAD boundaries is counter to findings from a recent cancer SV impact predictor(30) and may reflect StrVCTVRE’s focus on SVs that impact exons. Additionally, the low importance of deletion/duplication status suggests that on average, for exon-altering deletions and duplications, the region altered by an SV is more important than whether there was a gain or loss of genome content.

### Characterization of StrVCTVRE training and held-out test sets

A total of 7,263 pathogenic or likely pathogenic deletions and 4,551 pathogenic or likely pathogenic duplications were collected from ClinVar(31), a public database of variants cataloged by academic institutions and clinical laboratories. We restricted our data to deletions and duplications, as they are the only SV types with more than 500 pathogenic examples in ClinVar. Additionally, deletions and duplications constitute the vast majority (> 95%) of rare gene-altering SVs(6). A set of primarily benign SVs (described in greater detail below) were collected from ClinVar, gnomAD-SVs(16), and a recent great ape sequencing study(32). Because these ape SVs were mapped to the human genome, they may be biased towards more conserved genomic regions. We retained only rare (MAF < 1% in general population) SVs in order to match the challenge faced by SV discovery pipelines. Indeed, 92% of SVs identified through cohort sequencing are rare(16), so the salient challenge is to distinguish rare pathogenic SVs from rare benign SVs. Existing SV predictors have been trained and assessed on common benign SVs(23, 30), which may cause them to instead rely on features that separate common from rare SVs and result in lower accuracy in clinical use(26).

By training on rare SVs, we intend to achieve better accuracy in the challenge faced in pathogenic SV discovery. To create a rare benign dataset that matches the size range of our pathogenic dataset, we included SVs observed as homozygous at least once in great apes but rare in humans, which we assume should be mostly benign in humans due to our recent shared ancestry with great apes. Our benign dataset also included unlabeled rare SVs from gnomAD-SVs. Although we expect a small fraction of these unlabeled SVs are pathogenic, we made two assumptions that mitigated this issue: (1) pathogenic SVs have been depleted by selection so the large majority of unlabeled SVs are benign, and (2) the fraction of truly pathogenic SVs in the pathogenic and benign training sets is sufficiently different for StrVCTVRE to learn important distinguishing features. By including these additional data sources, we brought the ratio of pathogenic to benign SVs closer to 1:1 in our training set, even at small sizes. This would have been impossible with ClinVar data alone due to the dearth of small benign SVs in ClinVar.

To assess the appropriateness of including SVs from apes and gnomAD in our benign dataset, we explored how performance and feature importance changed with these data included. One predictor was trained only on ClinVar SVs, and a second predictor was trained on ClinVar SVs, ape SVs, and gnomAD SVs (altogether 3.8x more SVs than ClinVar alone). Using leave-one-chromosome-out cross validation, we found both training sets performed similarly (Fig. 2a), supporting our theory that the selected rare unlabeled gnomAD SVs and great ape SVs are sufficiently depleted in pathogenic SVs to be used as a training set of rare, benign SVs. Additionally, the predictor trained on all data showed a distribution of feature importance that is more evenly distributed among feature categories and possibly more robust. This includes a decrease in usefulness of gene importance features, which are likely to be overrepresented in ClinVar data, and an increase in importance in CDS features, which are an important line of evidence for assessing SV pathogenicity(33).

**Fig. 2.**
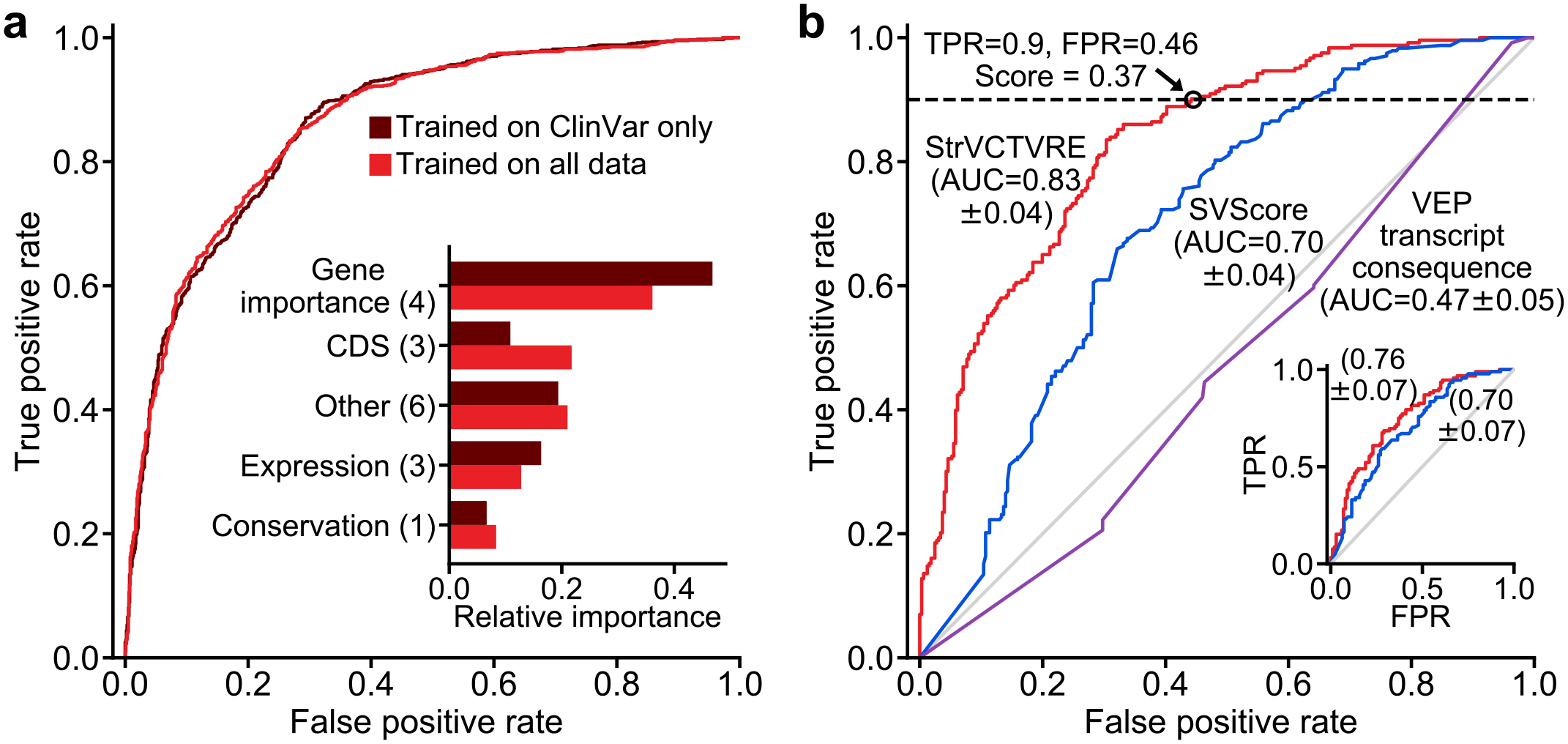
By training on multiple datasets, StrVCTVRE learned diverse feature importances and performed well on a held-out ClinVar test set. **a** Receiver operating characteristic comparing StrVCTVRE models trained on two different benign datasets: ClinVar in dark red, and all data (ClinVar, SVs common to apes but not humans, and rare gnomAD SVs) in medium red. When tested only on ClinVar data, performance does not significantly differ between the two training sets. However the feature importances (inset) of the classifier trained on all data (medium red) were more evenly distributed among feature categories. This suggests that unlabeled rare SVs and common ape SVs are a suitable benign training set. **b** Receiver operating characteristic comparing StrVCTVRE (red) to other methods on a held-out test set comprised of ClinVar SVs on chromosomes 1, 3, 5, and 7. Black circle indicates a StrVCTVRE score of 0.37, which we refer to as the ClinVar 90% sensitivity threshold. Inset shows performance on the same held-out test, modified so that each gene is overlapped by a maximum of 1 SV. AUC with 95% confidence interval is in parentheses.

Before training, all data were extensively cleansed to remove duplicate records within and between datasets, remove common SVs, and remove SVs larger than 3 Mb (see Methods). Pathogenic deletions and duplications were found to have a large size bias, likely due to the sensitivity of detection methods to specific size ranges (Fig. S2). To avoid training on this acquisition bias, putatively benign SVs were sampled to match the pathogenic SV size distribution (Fig. 3; Fig. S3). Specifically, in our training data we included only pairs of pathogenic and benign SVs that were of similar size and the same type (deletion or duplication). Using this matching strategy, we were able to include nearly all pathogenic deletions and duplications below 1 Mb. By incorporating ape and gnomAD SVs, we were able to include pathogenic SVs below 10 kilobases (kb), a range nearly absent in ClinVar benign SVs. In the benign training set, 26% of deletions and 75% of duplications came from ClinVar benign or likely benign SVs.

**Fig. 3.**
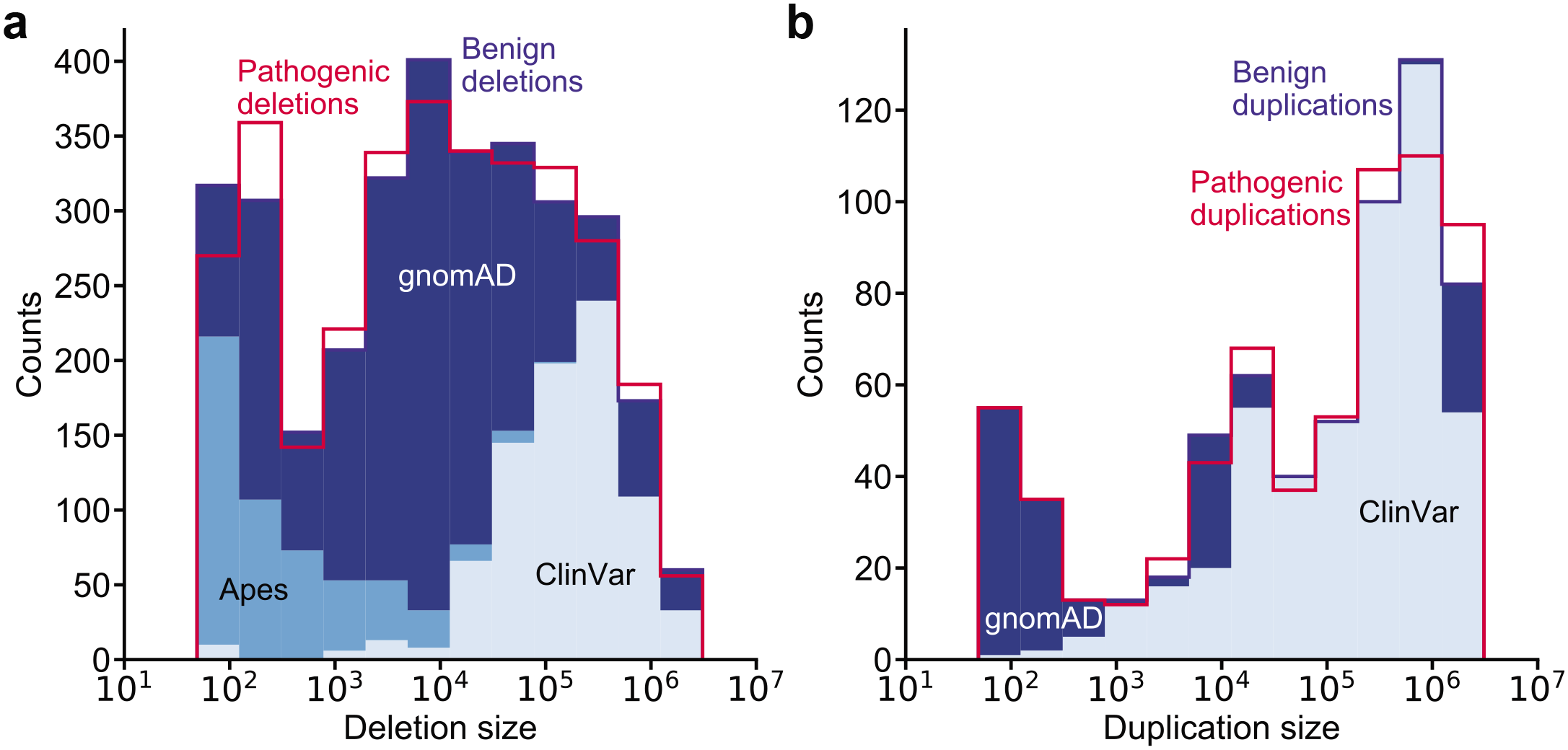
Benign training SVs (blue-shaded histograms) closely match the size distribution of pathogenic training SVs (red histogram outlines) and were drawn from multiple datasets. Histogram of pathogenic and benign (**a**) deletions and (**b**) duplications. **a** Benign deletions are composed of 26% ClinVar, 16% apes, and 58% gnomAD. **b** Benign duplications are composed of 75% ClinVar and 25% gnomAD. We were able to include more small pathogenic SVs in our training data by using apes and gnomAD SVs. Pathogenic SVs are composed entirely of ClinVar Pathogenic and Likely Pathogenic SVs and thus only histogram outlines are shown.

To accurately assess StrVCTVRE’s performance, we used a held-out test set of ClinVar SVs on chromosomes 1, 3, 5, and 7 (∼20% of the total ClinVar dataset). Only ClinVar SVs were used for testing since it is the highest-confidence dataset. The training set consisted of SVs from all three data sources on all remaining chromosomes. The training set consisted of 2,463 pathogenic SVs and 2,372 benign SVs, and the test set consisted of 244 pathogenic SVs and 334 benign SVs. The test set is of reduced size because pathogenic and benign SVs in the test set were matched on length. None of the SVs in the test set were used to develop the trained algorithm.

### StrVCTVRE eliminates more than half of benign SVs from consideration at 90% sensitivity

In discriminating between pathogenic and putatively benign ClinVar SVs in the test dataset, StrVCTVRE performed substantially better than published methods. Performance was measured using the area under the receiver operating characteristic curve (AUC). The AUC for StrVCTVRE was 0.83 (95% CI: 0.79 – 0.87). By comparison, SVScore had an AUC of 0.70 (95% CI: 0.66-0.74). StrVCTVRE improved notably in the classification of large duplications and deletions (> 1 MB), a regime in which SVScore by default classifies all SVs as pathogenic (lower left corner of Fig. 2b). We also evaluated the predictive ability of transcript consequence reported by VEP (AUC = 0.47; 95% CI: 0.42 – 0.52), and we found it performed no better than random. This poor performance was largely due to VEP annotating more benign SVs than pathogenic SVs with its most deleterious sequence ontology term, transcript ablation (Fig. S5). The poor performance of transcript consequence from VEP reinforces the known limitations of prioritizing variants using sequence ontology terms in isolation. As we intend StrVCTVRE to be used to prioritize SVs seen in clinical cases, it needs to perform well in clinically relevant regimes. Clinicians must limit cases in which pathogenic variants are misclassified as benign (false negatives), which requires strong performance at high sensitivity(34). When compared to existing methods, StrVCTVRE makes substantial improvements in the high-sensitivity regime, as it is able to capture 90% of pathogenic SVs at a 46% false positive rate (black circle, Fig. 2b). StrVCTVRE scores range from 0 to 1, with higher scores indicating a greater likelihood of pathogenicity. In Fig 2b, 90% sensitivity is reached at a StrVCTVRE score of 0.37, which suggests that when used on a collection of SVs called from a clinical cohort, this threshold may identify 90% of pathogenic SVs while reducing the candidate SV list by 54%. We refer to this StrVCTVRE score as the ClinVar 90% sensitivity threshold. StrVCTVRE performed equally well or better on test sets in which duplicates and common SVs were not removed or different size limits were imposed (Fig. S4).

We observed apparent clustering in the ClinVar data that led to additional analysis. Genes that are well-studied are overlapped by multiple pathogenic SVs catalogued in ClinVar. This resulted in several genes that were over-represented in our test set. Since SVs that overlap the same gene tend to be mostly pathogenic or mostly benign, this results in clustered test data, which may lead to higher variance in AUC performance. While this may yield improved performance for genes of particular interest, it may hide possible deficits in genome-wide performance. To address this, we randomly generated a test dataset in which each gene is overlapped by at most one SV (Fig. 2b inset). We found that the StrVCTVRE AUC was reduced when applied to this dataset, but StrVCTVRE was able to identify pathogenic SVs better than or equal to SVScore at all sensitivities. On this dataset, StrVCTVRE shows a sensitivity of 90% at a false positive rate of 59%.

### StrVCTVRE sensitivity threshold is validated on recent clinical SVs

To assess the accuracy of our ClinVar 90% sensitivity threshold and evaluate whether StrVCTVRE performs well on clinical data, we evaluated our method on a set of SVs identified by researchers at the Broad Institute Center for Mendelian Genomics (CMG). These SVs were recently identified through exome sequencing of patient cohorts with undiagnosed neuromuscular or retinal degeneration disorders(35–39). Clinical researchers determined these rare SVs were disease-causing or likely disease-causing. To avoid overlap between these CMG clinical SVs and StrVCTVRE training SVs, we used a leave-one-chromosome-out approach, in which 24 separate StrVCTVRE classifiers were developed, one for each chromosome. For example, CMG clinical SVs on chromosome 1 were predicted by a StrVCTVRE classifier trained on chromosomes 2, 3, 4, etc. The CMG clinical SVs consisted of 32 deletions and 2 duplications, were located on 14 chromosomes, and had a median size of 12kb (Table S1). At the ClinVar 90% sensitivity threshold (StrVCTVRE score >0.37), StrVCTVRE identified 31 of 34 disease-causing SVs (91%) as potentially pathogenic.

### Performance of StrVCTVRE on an independent test set from DECIPHER

All held-out test SVs, and a large fraction of training SVs, come from a single database: ClinVar. To independently test StrVCTVRE, we collected pathogenic and benign SVs from DECIPHER, a public database to which clinical scientists submit SVs seen in patients with developmental disorders(40). Because there is some overlap between training ClinVar SVs and DECIPHER SVs, we tested on DECIPHER using a leave-one-chromosome-out approach, as described above. Additionally, to ensure this DECIPHER test set is independent from our ClinVar test set, we considered only DECIPHER SVs with a reciprocal overlap of less than 10% with any SV used in training or testing StrVCTVRE. This strategy effectively removes any concerns of training and testing on the same or similar SVs. This test set included only DECIPHER variants with the highest classification confidence (Set 1, described below). Because StrVCTVRE was trained on SVs less than 3 Mb, and few benign SVs larger than 3 Mb have been observed(41), all SVs larger than 3 Mb were scored as pathogenic (given a score of 1). Compared to its performance on the ClinVar test set, StrVCTVRE performed similarly well on the DECIPHER test set, although performance varied across SV size (Fig. 4a). On large SVs (> 500 kb), StrVCTVRE performed very well (AUC = 0.91; 95% CI: 0.88 – 0.94; N=297), partially because most of the SVs larger than 3 Mb are correctly predicted as pathogenic. StrVCTVRE also performed very well (AUC = 0.89; 95% CI: 0.81 – 0.97, N=116) on small SVs (< 30 kb), although this is tempered somewhat by the relatively few small SVs in the DECIPHER dataset. StrVCTVRE performed well (AUC = 0.80; 95% CI: 0.72 – 0.88,N=545) on mid-length SVs, identifying pathogenic SVs significantly better than SVScore.

**Fig. 4.**
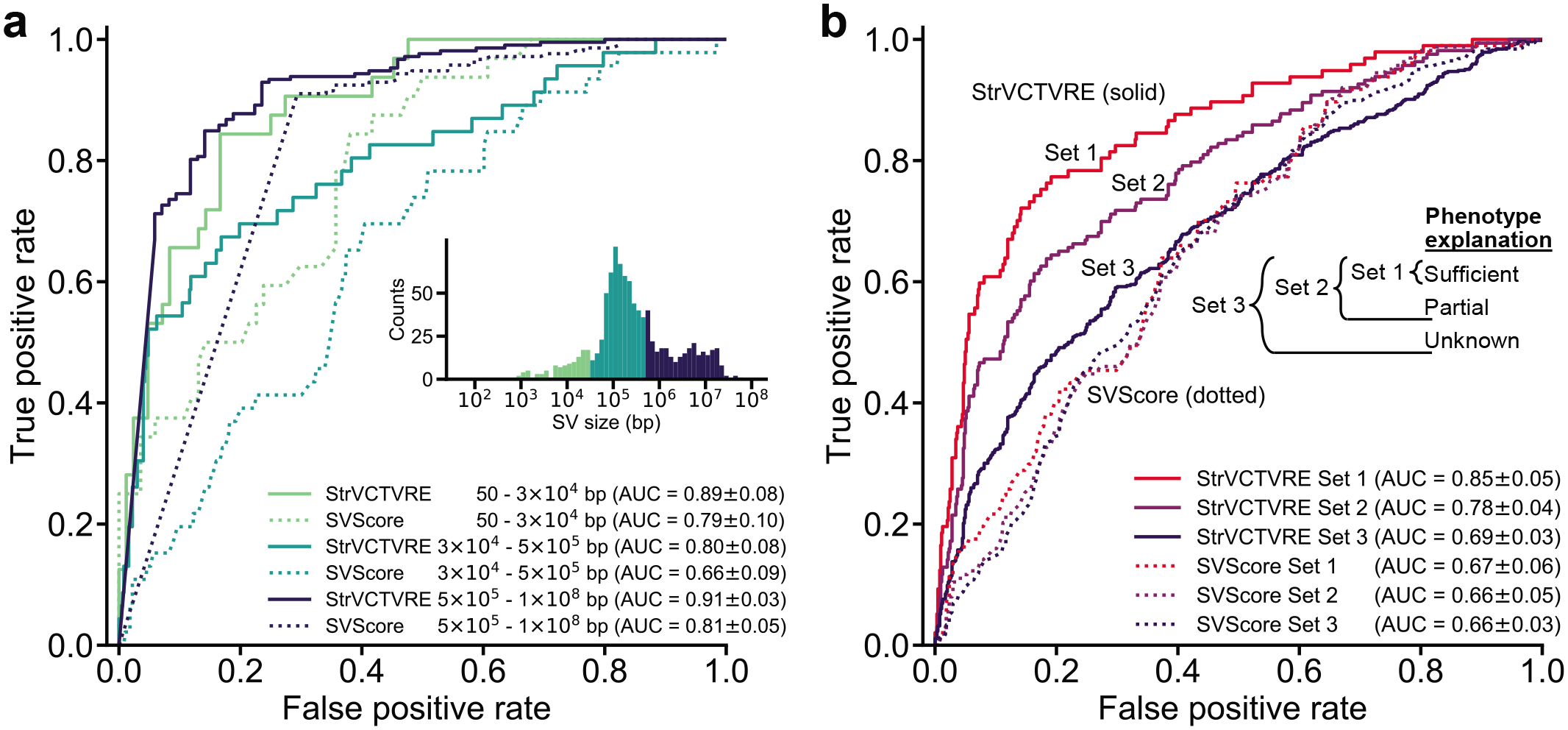
**a** Across three size ranges, StrVCTVRE accurately classified variants in an independent test set. In this ROC comparison of StrVCTVRE (solid line) and SVScore (dotted line), three size ranges of SVs were considered. StrVCTVRE performed very well on large and small SVs, while performing well on mid-sized variants. **b** When presented with data that are more reliably classified, StrVCTVRE’s performance improved. ROC plot showing StrVCTVRE’s performance increased as SV contribution to proband phenotype increases from set 3 (includes less confidently classified SVs) to set 2 and from set 2 to set 1 (most confidently classified SVs). The performance of SVScore did not significantly differ between the sets.

### StrVCTVRE performance is higher when assessed on more reliably classified data

We expect that some DECIPHER pathogenic SVs are in reality benign. SVs that better explain patient phenotype are more likely to be pathogenic. To investigate the effect of SV pathogenicity on predictor performance, we grouped DECIPHER SVs into 3 sets. Set 1 consisted of SVs that sufficiently explain the proband phenotype, and these should be reliably pathogenic. Set 2 included SVs that partially explain the proband phenotype and Set 1 SVs. Set 3 included SVs with unknown contribution to proband phenotype and Set 2 SVs, and therefore their pathogenicity is less certain. StrVCTVRE was tested using a leave-one-chromosome-out approach, and DECIPHER SVs were filtered based on overlap with training and testing data as described above. We found a consistent trend towards more accurate StrVCTVRE classification in sets that were more enriched for pathogenic SVs (Fig. 4b). However, the same trend was not observed for SVScore. Since StrVCTVRE’s performance improves on presumably more reliably classified data, we have reason to believe StrVCTVRE is making meaningful classifications.

### StrVCTVRE eliminates the most benign SVs seen in 221 individuals

Typically, patients with a rare disorder caused by homozygous SVs have one or two pathogenic SVs in their genome, and the remaining SVs are benign. An ideal impact predictor would prioritize the pathogenic homozygous SVs and eliminate from consideration as many of the benign SVs as possible. To evaluate StrVCTVRE’s performance in this scenario, we applied it to SVs called in 2,504 genomes identified by the 1000 Genomes Project phase 3(42) (1KGP). Because 1KGP should be depleted in individuals with severe rare disorders, we treated each genome as if it came from a proband with a rare disorder whose pathogenic SVs have been removed. 221 of these genomes had 1 or more homozygous rare exon-altering SVs, almost all of which should be benign. For each genome, we recorded the fraction of putatively benign SVs that were correctly identified as benign by StrVCTVRE and SVScore (Fig. 5a). Since many genomes had just one homozygous exon-altering SV, the distribution is bimodal at 0 and 1. We used our leave-one-chromosome-out predictors (e.g., predicting on 1KGP SVs on chromosome 1 and training StrVCTVRE on all other chromosomes) to score each SV. At the ClinVar 90% sensitivity threshold (StrVCTVRE score >0.37), on average StrVCTVRE identified 59% of the putatively benign SVs in each genome as benign, compared to 43% when SVScore was used at the same sensitivity (Wilcoxon paired-rank p = 8.06e-6). In a clinical setting, StrVCTVRE may classify more benign SVs as benign than SVScore, allowing clinicians and researchers to eliminate the most benign homozygous SVs from consideration.

**Fig. 5.**
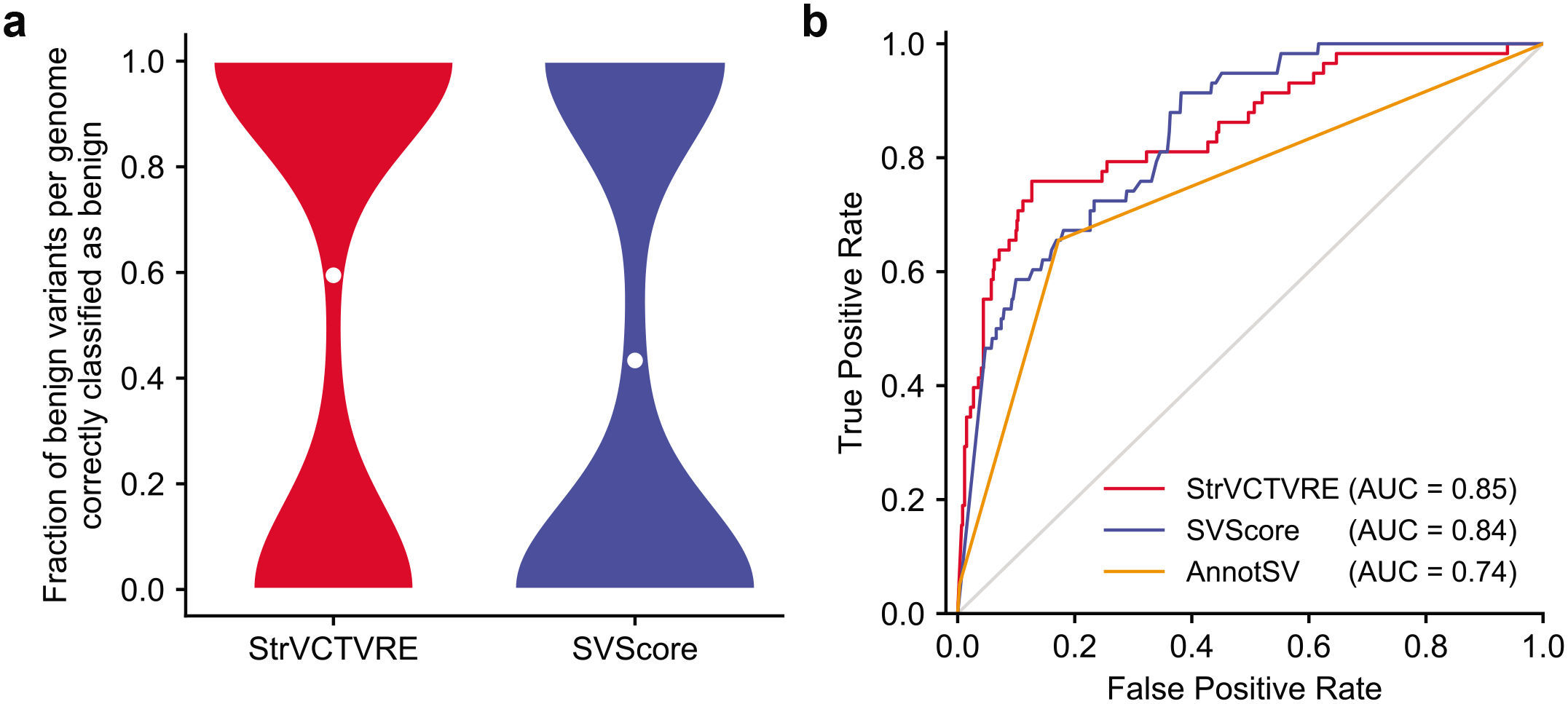
**a** StrVCTVRE eliminated a significantly larger fraction of benign SVs from consideration than SVScore. When tested on rare exonic SVs from the genomes of 221 putatively healthy individuals, StrVCTVRE was able to correctly classify 59% of putatively benign variants in each genome. White dots represent mean values. For both methods, the threshold for variant consideration was at the ClinVar 90% sensitivity (Fig. 2b). **b** ROC comparing two machine-learning methods with diverse features (StrVCTVRE and SVScore) to one method (AnnotSV) that uses limited features and manually determined decision boundaries. AnnotSV ranks an SV as ‘pathogenic’ or ‘likely pathogenic’ when the SV overlaps a catalogued pathogenic SV, known disease gene, or gene predicted to be intolerant to variation. To generate this figure, all SVs overlapping any of AnnotSV’s catalogued pathogenic SVs were removed from the DECIPHER Set 3 dataset, and the remaining SVs were used for testing. AnnotSV performs relatively poorly on these novel variants. In contrast, the machine learning methods perform better, possibly because they use more diverse features and have decision boundaries trained on real data. StrVCTVRE scores were generated using a leave-one-chromosome-out approach.

### StrVCTVRE performance is reliable even on SVs that do not overlap cataloged pathogenic SVs

Since probands with the same disorder often have SVs altering the same genome element, and recurrent pathogenic de novo SVs are known to occur(43), one strategy used to prioritize SVs is to annotate them with overlapping SVs of known pathogenicity. AnnotSV is a popular method to identify pathogenic SVs based on their overlap with both cataloged pathogenic SVs in the National Center for Biotechnology Information’s dbVar. Because it considers catalogued SVs, AnnotSV would likely perform very well for a proband whose disease-causing SV overlaps a cataloged pathogenic dbVar SV (Fig. S6). Yet, many probands have disease-causing SVs that are not cataloged. To address these novel SVs, AnnotSV also considers SV overlap with genes associated with disease or predicted to be intolerant to variation, and it uses manually determined decision boundaries to score SVs (e.g., an SV overlapping a gene with pLI > 0.9 is scored as likely pathogenic). To compare the performance of AnnotSV with machine learning SV impact predictors on novel SVs, we created a dataset of Set 3 DECIPHER SVs that do not overlap dbVar SVs used by AnnotSV, and we recorded the prediction accuracy of each method (Fig. 5b). AnnotSV performed notably worse on these uncatalogued SVs. We tested StrVCTVRE (using the leave-one-chromosome-out approach) and SVScore on these uncatalogued SVs, and both methods showed significant predictive power, which we attribute to their consideration of features beyond gene intolerance (such as conservation and expression features) and their use of methods that learn decision boundaries based on training data, rather than manually determined boundaries.

### Interpreting StrVCTVRE scores

StrVCTVRE scores range from 0 to 1, reflecting the proportion of decision trees in the random forest that classify an SV as pathogenic. Note that StrVCTVRE scores are not probabilities. Although we used the ClinVar 90% sensitivity threshold for evaluation, we advise against using StrVCTVRE scores as a threshold. We instead recommend that greater consideration be given to SVs with greater StrVCTVRE scores. However, thresholds are currently required for computational tools when SVs are classified using the guidelines for sequence variant interpretation recommended by the American College of Medical Genetics and Genomics (ACMG; criteria PP3, BP4)(33, 44). Within the ACMG framework, StrVCTVRE can be used as supporting evidence since it uses multiple lines of computational data. We suspect that higher levels of evidence (e.g., moderate) may be achievable, as shown by Tavtigian et al.(45) However, when using StrVCTVRE at higher levels of evidence, users should be careful not to also count other ACMG criteria that StrVCTVRE already incorporates, which could lead to double counting. Alternatively, to resolve concerns of double counting, StrVCTVRE can be used just to prioritize variants, but not used as evidence. Users then can manually classify SVs of interest using the full ACMG criteria.

## Discussion

As genome sequencing becomes more accessible, clinicians and researchers face a challenge in identifying pathogenic SVs in the thousands identified by sequencing. The ACMG recently offered guidelines for classifying SVs, acknowledging that classification is complex and many pathogenic SVs will be classified as variants of uncertain significance due to incomplete knowledge(33). SV impact predictors can address this challenge, but few SV impact predictors exist. Although SVs comprise a significant fraction of the loss-of-function mutations that cause rare disease, fewer than 10,000 pathogenic SVs have been cataloged in ClinVar. These SVs have distinct biases towards certain genes and lengths, which leads to acquisition bias that hinders predictor development. Additionally, it is not clear which features are most useful when classifying SVs and how to address the large size range of SVs. StrVCTVRE is the first method to address these problems by predicting the impact of exon-altering deletions and duplications in rare genetic disorders. We overcame data limitations and bias by combining SVs from multiple data sources as well as matching pathogenic and benign SVs by size. Since clinicians and researchers must recognize SVs that cause disease among dozens of rare exon-altering SVs detected in a proband, we trained only on rare SVs.

Determining whether a single SV is pathogenic requires consideration of numerous features in combination, as demonstrated by the recent ACMG SV guidelines. Independent of these guidelines, our method identified important features in cataloged SVs. Our findings reinforce clinical guidelines, while also highlighting new areas to explore. Both StrVCTVRE and the ACMG guidelines found gene importance and CDS disruptions to be critical for SV interpretation. Additionally, StrVCTVRE highlighted two features not discussed in the guidelines: conservation and expression. We found exon expression in particular is both predictive and poorly correlated with all other features, suggesting it captures distinctive information for determining pathogenicity. More widespread consideration of expression features could be beneficial for SV classification. StrVCTVRE additionally identified features that are not useful to classify exon-altering SVs, such as TAD boundary strength and whether there is a copy gain or loss. This is consistent with the ACMG guidelines, which do not consider TAD boundaries and provide very similar scoring metrics for both copy gain and loss.

Since SVs range from 50 bp to > 10 Mb, it is challenging to accurately classify SVs across this range. Benign SVs in ClinVar are mainly > 10 kb, but accurate classification of SVs < 10 kb requires training on benign SVs from the same size range. We accomplished this by training on small benign SVs from great apes and gnomAD. When tested on an independent test set, StrVCTVRE performed well at all size ranges. To be helpful in a clinical setting, a method must perform well at moderately high sensitivity. StrVCTVRE satisfies this requirement and was able to remove 57% of homozygous SVs from consideration at a sensitivity of 90% in the 1KGP dataset. This 90% sensitivity threshold was validated using a dataset of recent SVs observed to cause neuromuscular and retinal degeneration disorders. Overall, we found StrVCTVRE outperforms SVScore in most tasks, even though SVScore’s underlying approach, CADD, was trained on > 1,000-fold more variants. Additionally, whereas StrVCTVRE was often assessed using a leave-one-chromosome-out approach, SVScore could not be readily modified and thus had the benefit of possibly training on data that overlapped the testing SVs.

StrVCTVRE is accessible as a downloadable command line program (see Data Availability). Whereas SVScore requires users to download an 80 gigabyte (Gb) CADD file, StrVCTVRE only requires a 9 Gb phyloP file. Because there are an intractably large number of possible SVs, each SV must be scored anew (unlike SNVs for which scores can be pre-computed), and this requires efficient scoring methods. StrVCTVRE runs rapidly and annotates 100,000 gnomAD SVs in three minutes, while SVScore annotates the same SVs in 24 hours (Fig. S7).

Following existing predictors, StrVCTVRE predicts the pathogenicity of an SV in isolation. Yet human biology complicates this picture through zygosity and dominance. Since zygosity is not reported for most SVs in ClinVar, StrVCTVRE is zygosity-naïve. Additionally, StrVCTVRE’s pathogenic training dataset consists largely of SVs in genes predicted to lead to dominant disorders (Fig. S8). When tested on sets of predicted dominant or recessive SVs, StrVCTVRE performs similarly on both (Fig. S9). Researchers who suspect a recessive mode of inheritance may need to consider StrVCTVRE scores in tandem with impact predictor scores for SNVs in trans in the same gene. Although genes vary in their tolerance of SVs and dominance, we believe a whole genome approach will be necessary to identify all pathogenic SVs, including those SVs disrupting genes not currently associated with disease. To identify new disease genes, it may be helpful to consider StrVCTVRE scores in tandem with one of the many methods that assess the match between patient phenotype and known/predicted phenotypes for an affected gene(46–48).

A method can only be as good as its training data. SV impact predictors are limited by the relatively small number of identified pathogenic and putatively benign SVs, as well as the over-representation of certain genes in the dataset (Fig. S1). While pathogenic ClinVar variants are commonly used to train variant impact predictors, they are known to include misclassified variants(49). We know of no characterization of the accuracy of SVs in ClinVar, but work investigating pathogenic SNVs suggest at least 90% are pathogenic based on reclassification rates(50). 70% of our pathogenic training SVs have at least 1 review star in ClinVar, indicating they have supporting evidence which further bolsters our confidence in these data.

Nonetheless, data limitations almost certainly curtail the ultimate performance of our approach. StrVCTVRE is unable to classify inversions and insertions due to limited data; however, these have been shown to contribute to a minority of the pLoF events caused by SVs(16). We are hopeful that additional clinical sequencing studies will identify a more diverse range of SVs, which will be cataloged in open resources such as ClinVar and leveraged to develop more accurate models. We look forward to greater non-coding genome annotations, which will expand our understanding and cataloging of pathogenic noncoding SVs, which remain vexing to classify.

## Conclusions

Much of the focus in SV algorithms has been on methods to accurately detect SVs. These methods have left clinicians and researchers awash with SVs not previously known. As experimental methods and algorithms advance, SV detection will improve, but SV interpretation will continue to be challenging. StrVCTVRE advances the clinical evaluation of SVs. During genome sequencing analysis, some cases contain an SV that matches a cataloged pathogenic SV or satisfies the conditions for pathogenicity set forth in the ACMG SV guidelines. However, these SVs are often not obvious, and StrVCTVRE can be used to quickly bring these SVs to attention. In the many cases in which no SV is immediately promising, StrVCTVRE aids clinicians and researchers in identifying compelling SVs for manual investigation. Then, if a case remains unresolved by manual investigation, SVs highlighted by StrVCTVRE that are in novel disease genes can be directed to experimental exploration. This will empower researchers to identify novel disease genes where haploinsufficiency and triplosensitivity were not previously known causes of disease. Adoption of structural variant impact predictors will enable clinicians and researchers to make the most of these new data to improve both patient care and our understanding of basic biology.

## Methods

### Training, validation, and test datasets

All SVs were retrieved in GRCh38 or converted using the University of California, Santa Cruz (UCSC) liftover tool(51).

All ClinVar SVs(31) were downloaded from ftp://ftp.ncbi.nlm.nih.gov/pub/clinvar/tab_delimited/variant_summary.txt.gz on January 21, 2020. SVs were retained if they fulfilled all the following requirements: clinical significance of pathogenic, likely pathogenic, pathogenic/likely pathogenic, benign, likely benign, or benign/likely benign; not somatic in origin; type of copy number loss, copy number gain, deletion, or duplication; > 49 bp in size; at least 1 bp overlap with an exon.

Great ape SVs(32) mapped to GRCh38 were downloaded from ftp://ftp.ebi.ac.uk/pub/databases/dgva/estd235_Kronenberg_et_al_2017/vcf/ on April 8, 2019. Deletions were retained if they were absent in humans and homozygous in exactly one of the following species: chimpanzee, gorilla, or orangutan. Only exon-altering deletions > 49 bp were retained. These deletions are subsequently referred to as *apes*.

gnomAD 2.1.1 SVs(16) (build GRCh37) were downloaded from https://storage.googleapis.com/gnomad-public/papers/2019-sv/gnomad_v2.1_sv.sites.vcf.gz on June 28, 2019. Only duplications and deletions were retained that were exon-altering, >49 bp, and PASS Filter. gnomAD SVs were divided into three categories: SVs with a global minor allele frequency (MAF) > 1% (*gnomAD common*), SVs with a global MAF < 1% with at least one individual homozygous for the minor allele (*gnomAD rare benign*), and SVs with a global MAF < 1% with no individuals homozygous for the minor allele (*gnomAD rare unlabeled*).

Database of Genomic Variants(41) release 2016-05-15 of GRCh38 “DGV Variants” was downloaded from http://dgv.tcag.ca/dgv/app/downloads on April 08, 2019. MAF of each deletion was calculated as ‘observedlosses’ / (2 * ‘samplesize’). MAF of each duplication was calculated as ‘observedgains’ / (2 * ‘samplesize’). Only exon-altering SVs > 49 bp were retained. Those SVs with a MAF greater than 1% are subsequently referred to as *DGV common*.

DECIPHER CNVs (build GRCh37) were downloaded from http://sftpsrv.sanger.ac.uk/ on Jan 27, 2020. Only exon-altering SVs > 49 bp with pathogenicity of “pathogenic”, “likely pathogenic”, “benign”, or “likely benign” were retained. We only considered benign or likely benign SVs without “Full” or “Partial” contribution to disease phenotype. These benign and likely benign SVs were included in all 3 sets. Set 1 pathogenic SVs consisted of pathogenic or likely pathogenic SVs with “Full” contribution to disease phenotype (referred to as “sufficient” in this manuscript). Set 2 SVs consisted of pathogenic or likely pathogenic SVs with “Full” or “Partial” contribution. Set 3 SVs consisted of pathogenic or likely pathogenic SVs with “Full”, “Partial”, or “Unknown” contribution. Identical SVs with conflicting pathogenicity were removed. SVs were then sorted by size (ascending) and SVs with a reciprocal overlap >90% were removed, keeping only the first SV.

1KGP merged SVs(42) were downloaded from ftp://ftp.1000genomes.ebi.ac.uk/vol1/ftp/phase3/integrated_sv_map/supporting/GRCh38_positio ns/ on Oct 22, 2019. Only exon-altering deletions and duplications with a global allele frequency less than 1% were used for testing in Fig. 5a.

We used exon boundaries from Ensembl biomart(52), genes v96, GRCh38.p12, limited to genes with HGNC Symbol ID(s) and APPRIS annotation(28). For each gene, a single principal transcript was used, based on the highest APPRIS annotation. For transcripts that tied for highest APPRIS annotation, the longest transcript was used. Exon overlap was determined using bedtools intersect.

Extensive deduplication of data was performed as follows. Deletions and duplications were considered separately. Benign SVs (n=23,239) were ordered (ClinVar benign, ClinVar likely benign, apes, gnomAD rare benign, gnomAD rare unlabeled) and duplicates (reciprocal overlap of 90% or greater) were removed, keeping the first appearance of an SV. This removed 577 SVs from ClinVar benign/likely benign, 5 SVs from apes, and 408 SVs from gnomAD. The retained data are subsequently referred to as *benign.* To deduplicate pathogenic SVs (n=8,378), deletions and duplications were considered separately. Exact matches between ClinVar pathogenic and ClinVar likely pathogenic were removed from likely pathogenic. SVs were then sorted by size, ascending. SVs with > 90% reciprocal overlap were removed, keeping the smallest SV. This removed 2,421 pathogenic SVs. The retained data are subsequently referred to as *pathogenic.* Next, exact matches between the benign and pathogenic datasets were removed from both datasets. Finally, duplicates between pathogenic and benign (reciprocal overlap of 90% or greater) were removed from the pathogenic dataset. This removed 3 benign SVs and 82 pathogenic SVs.

Data were processed as follows to ensure we trained only on rare SVs. Pathogenic and benign SVs that exactly matched a DGV common SV were removed. Pathogenic and benign SVs with reciprocal overlap > 90% with an SV in gnomAD common were removed. This removed 30 benign SVs and 1 pathogenic SV. SVs between 50 bp and 3 Mb were retained, all others were removed.

We found some evidence of acquisition bias in ClinVar data due to the SV size sensitivity of different SV detection methods (Fig. S2). To ensure StrVCTVRE was not learning on this acquisition bias, the size distribution of benign and pathogenic SVs were matched using the following procedure. After filtering as described above, benign SVs were organized into five tiers: ClinVar likely benign; ClinVar benign; apes; gnomAD rare benign; and gnomAD rare unlabeled. Each pathogenic SV was then matched by size and type (DEL or DUP) to a benign SV, iterating through each tier. Specifically, each pathogenic SV of size *N* seeks a benign SV of the same type in the bin 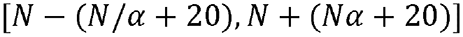 where 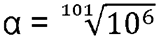 (this bin size derived from Ganel et al.(23)). A pathogenic SV first seeks a benign SV in the first benign tier. If matched, the pathogenic and benign SVs are included in the training set, and the benign SV cannot match any further pathogenic SVs. If no match is found in the first benign tier, the same process is repeated while progressing through further benign tiers. Pathogenic SVs that do not find a match in any benign tier are not included in the final training set. This process was continued for all pathogenic SVs and the resulting data are shown in Fig. 3 and S3.

After SVs were annotated with features (see below), we identified groups of SVs with identical features, considering pathogenic and benign SVs separately. We removed all but one of these feature-identical SVs in order to avoid overfitting. This removed 37 SVs from the pathogenic training set and 31 SVs from the benign training set. For feature-identical SVs that were present in both the pathogenic and the benign datasets, all feature-identical SVs were removed. This removed 13 SVs.

### Structural variant impact predictors

VEP(17) v96 was downloaded from https://github.com/Ensembl/ensembl-vep on April 16, 2019, and used to annotate SVs with transcript consequence sequence ontology terms. SVScore(23) v0.6 was downloaded from https://github.com/lganel/SVScore on June 16, 2019. It was run using CADD(24) v1.3, downloaded from https://cadd.gs.washington.edu/download on June 16, 2019, using default settings. AnnotSV(20) v2.3.2 was downloaded from https://github.com/lgmgeo/AnnotSV on Feb 27, 2020. AnnotSV was run using human annotation and default settings.

### Structural variant features

All gene and exon boundaries used to determine features came from Ensembl Genes v96 as described above. Each SV was annotated with the following 17 features:

**Table 2:**
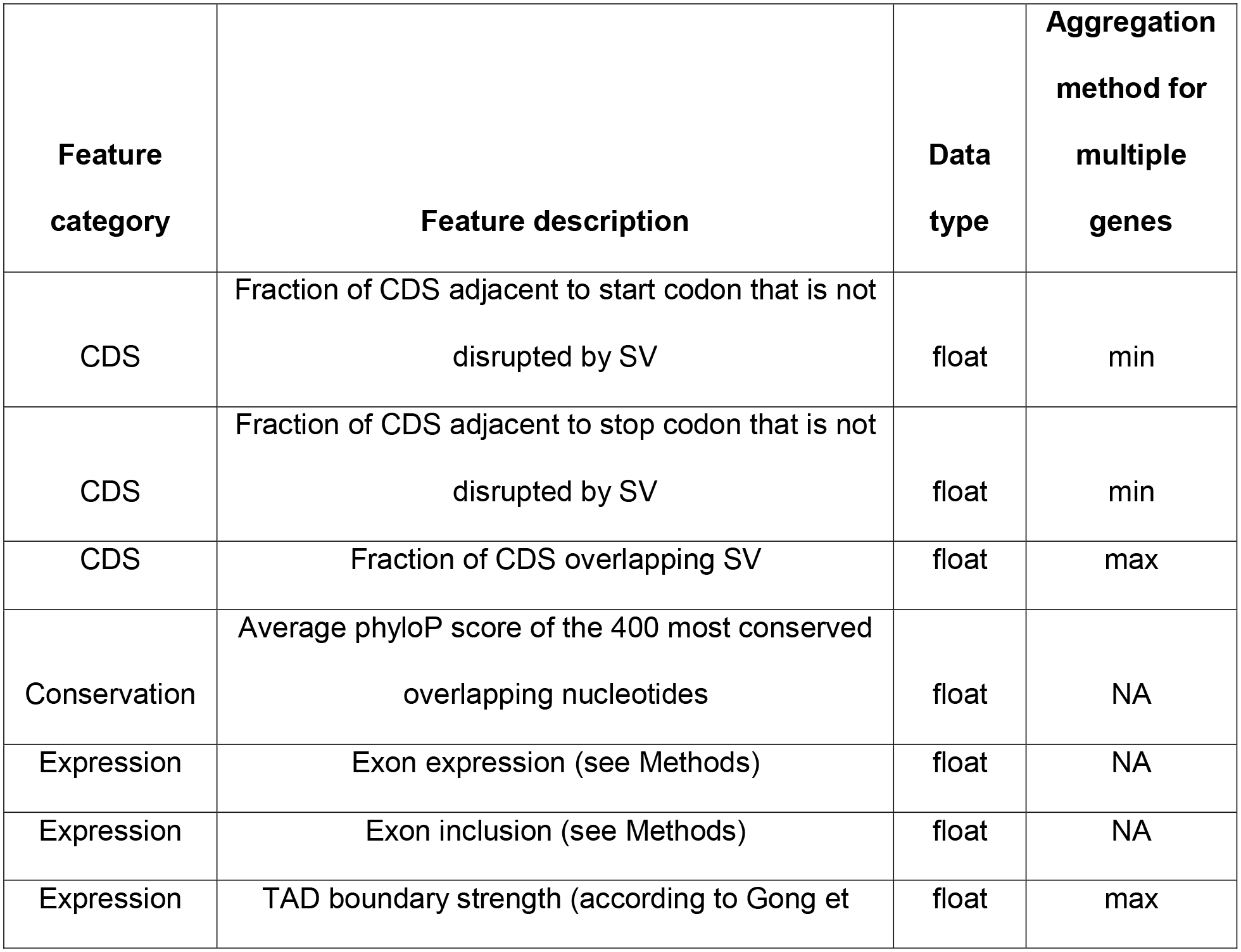
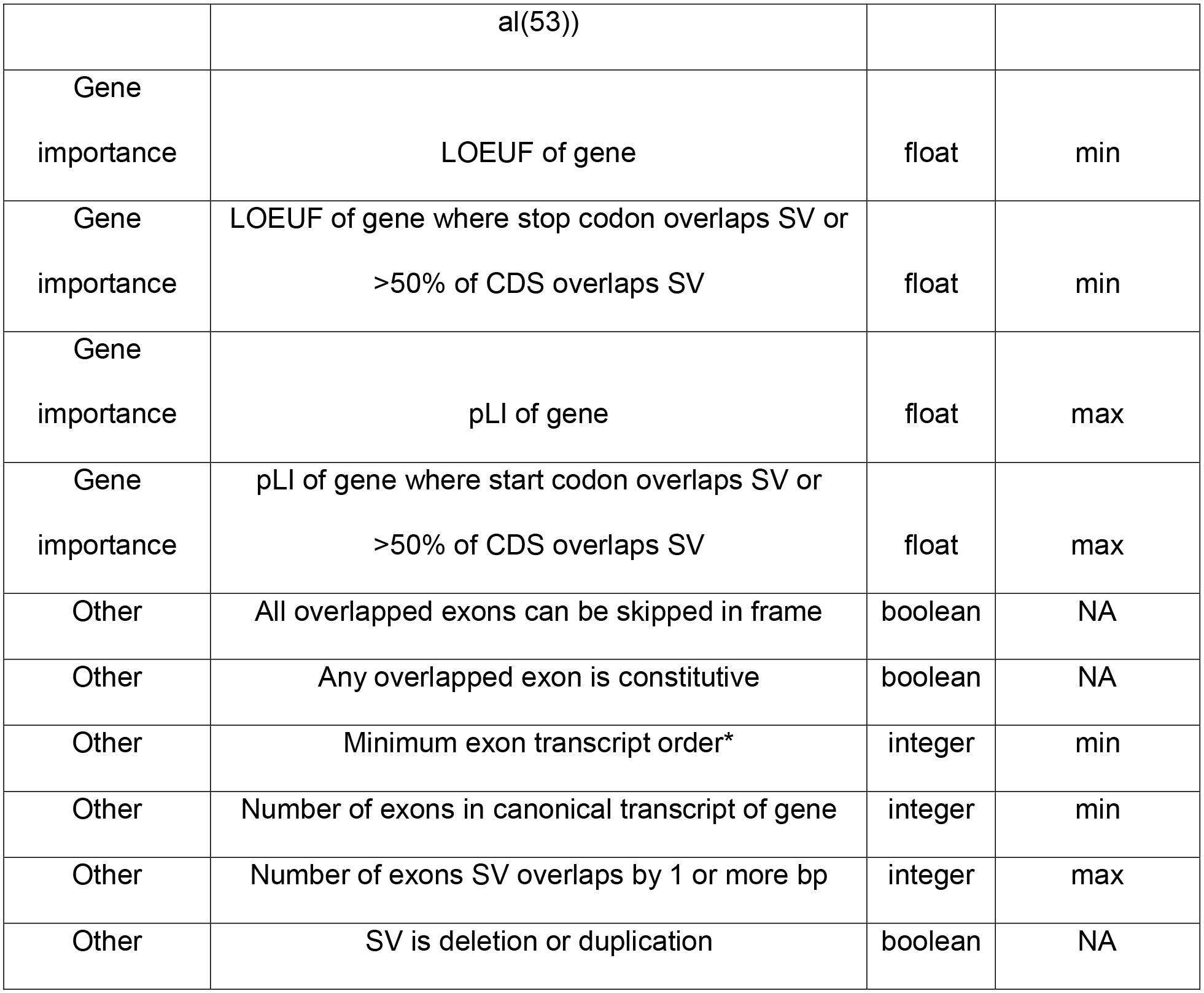
Features used in StrVCTVRE. *exon transcript order was defined as the number of exons preceding a given exon in a gene.

| Feature category | Feature description | Data type | Aggregation method for multiple genes |
| --- | --- | --- | --- |
| CDS | Fraction of CDS adjacent to start codon that is not disrupted by SV | float | min |
| CDS | Fraction of CDS adjacent to stop codon that is not disrupted by SV | float | min |
| CDS | Fraction of CDS overlapping SV | float | max |
| Conservation | Average phyloP score of the 400 most conserved overlapping nucleotides | float | NA |
| Expression | Exon expression (see Methods) | float | NA |
| Expression | Exon inclusion (see Methods) | float | NA |
| Expression | TAD boundary strength (according to Gong et al(53)) | float | max |
| Gene importance | LOEUF of gene | float | min |
| Gene importance | LOEUF of gene where stop codon overlaps SV or >50% of CDS overlaps SV | float | min |
| Gene importance | pLI of gene | float | max |
| Gene importance | pLI of gene where start codon overlaps SV or >50% of CDS overlaps SV | float | max |
| Other | All overlapped exons can be skipped in frame | boolean | NA |
| Other | Any overlapped exon is constitutive | boolean | NA |
| Other | Minimum exon transcript order* | integer | min |
| Other | Number of exons in canonical transcript of gene | integer | min |
| Other | Number of exons SV overlaps by 1 or more bp | integer | max |
| Other | SV is deletion or duplication | boolean | NA |

Expression features were derived from transcript data downloaded from the GTEx Portal v7(54). Exon expression was calculated for each nucleotide as the sum of the transcripts per million (TPM) of fragments that map to that nucleotide. Exon inclusion estimated the proportion of transcripts generated by a gene that include a given nucleotide and was calculated for each nucleotide as the TPM of fragments that map to that nucleotide, divided by the sum of TPM that map to the gene containing that nucleotide. For both features, adjacent base pairs with the same value were merged together into genomic intervals. For SVs that overlapped more than one of these genomic intervals, exon expression was calculated by averaging the 400 highest exon expression genomic intervals contained in that SV. The same was done for exon inclusion. All GTEx tissues were used in this analysis.

To determine which conservation feature to use, we assessed the accuracy of both PhastCons(55) and PhyloP(29) in discriminating between pathogenic and benign SVs using the average of the highest-scoring 200, 400, 600, 800, and 1000 nucleotides (Fig. S10). The test set consisted of 200 small (< 800 bp) SVs randomly selected from our pathogenic and benign SV training datasets (as described above). We found the mean PhyloP score of the 400 most conserved nucleotides in an SV was among the highest accuracy predictors. For both conservation and expression features, if the total overlap between the SV and all exons was less than 400 intervals, then the values of the overlapped intervals were averaged together to calculate the feature. Median imputation was used to fill in missing feature annotations.

In Fig. 1a, features were clustered by correlation using the linkage and fcluster functions from the SciPy(56) v 1.1.0 hierarchical clustering package. The input to this figure were the features for all SVs used as training data. Values for some features were reversed to ensure most matrix correlations are positive.

### Random forest classification

StrVCTVRE was implemented as a random forest classifier in Python with scikit-learn(57) v0.17, using class RandomForestClassifier. A grid search was performed to find the optimal hyperparameters by using a leave-one-chromosome-out cross validation strategy and validation only on ClinVar data, as described previously. The hyperparameters searched included: the max depth of a tree (5, 10, 15, No limit), max features considered at each split (1, 2, 3, 4), the minimum samples at each leaf node (1, 2, 4), the minimum samples required to split a node (2, 4), the number of trees generated (500, 1000, 3000), and whether to use out-of-bag samples to estimate accuracy (True, False). Several combinations of features performed similarly well, and we chose one that performed well while unlikely to over-fit to the training data—max depth: 10, max features considered at each split: 1, minimum samples at each leaf node: 2, minimum samples required to split a node: 4, number of trees: 1,000, out of bag samples: False. Feature importance used in figures is also known as Gini importance(58), and was calculated using the feature_importances_ attribute of RandomForestClassifier.

### Figures

In Fig. 1b, 95% confidence intervals were derived by generating 1,000 random forest predictors.

In Fig. 2a, the data were generated by using a leave-one-chromosome out approach that included all chromosomes besides chromosomes 1, 3, 5, and 7 (e.g., SVs in chromosome 2 were assessed using training data from chromosomes 4, 6, 8, 9, 10, etc.).

In Fig. 2b, to create the inset testing set, we began with the benign and pathogenic datasets as described above, and only retained ClinVar SVs from each dataset. Next, we removed any SVs larger than 3MB, and for both the benign and pathogenic dataset, we randomly sampled SVs without replacement, such that SVs were retained if they did not overlap any of the same genes as a previously sampled SV. This resulted in a reduced dataset for both pathogenic and benign SVs, in which every gene was overlapped by at most a single SV. Pathogenic and benign SVs from these reduced datasets were then matched by size as described above, and only results from testing on SVs on chromosomes 1, 3, 5, and 7 are shown in the Fig. 2b inset.

In Fig. 2b, 4a, and 4b, AUC 95% confidence intervals were derived by calculating the AUC standard error following Hanley and McNeil(59).

In Fig. 4b, 90% sensitivity thresholds were derived from StrVCTVRE and SVScore performance on the ClinVar held-out test set (dotted line, Fig. 2b).

### Method availability

The StrVCTVRE command line tool can be downloaded from https://compbio.berkeley.edu/proj/strvctvre.

## Declarations

### Ethics approval and consent to participate

Not applicable

### Consent for publication

Not applicable

### Availability of data and materials

All datasets generated and analyzed during the current study are available in the Dryad repository, https://datadryad.org/stash/share/i6z9fpHWtFW7rzoT9Zv8yMG5eq15pdltndY6aJwMCik, with the following exceptions. Additional data were obtained from DECIPHER, for which access was granted for the current study, but these data are not publicly available due to their sensitive nature. Under reasonable request, DECIPHER data can be requested from https://www.deciphergenomics.org/about/data-sharing. A subset of the recent CMG clinical SVs found to cause neuromuscular and retinal degeneration disorders have been made publicly available, but the full dataset is not yet publicly available due to the sensitive nature of the data. These data can be made available by Dr. Anne O’Donnell on reasonable request.

### Authors’ contributions

AGS designed and assessed the method. ZH performed additional analyses. AGS, ZH, and SEB wrote the manuscript. SEB and SRS supervised the work. All authors revised the manuscript. All authors have read and approved the final manuscript.

## Acknowledgements

We thank Dr. Anne O’Donnell, Dr. Julia Goodrich, Grace Tiao, Katherine Chao, and Isaac Wong for providing rare clinical SVs. We thank Dr. Nilah Ioannidis for helpful feedback during the project. We thank Dr. Aashish Adhikari for insightful comments in the initial planning of the project. We thank Azza Althagafi for thorough testing of our GitHub resources, as well as Lindsey Guan, Reet Mishra, and Ashish Ramesh for early script testing. We thank Drs. Véronique Geoffroy and Jean Muller for helpful discussion of an earlier preprint. This study makes use of data generated by the DECIPHER Consortium. A full list of centers who contributed to the generation of the data is available from http://decipher.sanger.ac.uk and via email from. SEB was unable to review this manuscript in its entirety due to injury. A preprint of this work was made available at https://doi.org/10.1101/2020.05.15.097048 on May 16, 2020.

## Corresponding author

Steven E. Brenner

