## Supplemental Figures for "StrVCTVRE: A supervised learning method to predict the pathogenicity of human genome structural variants"

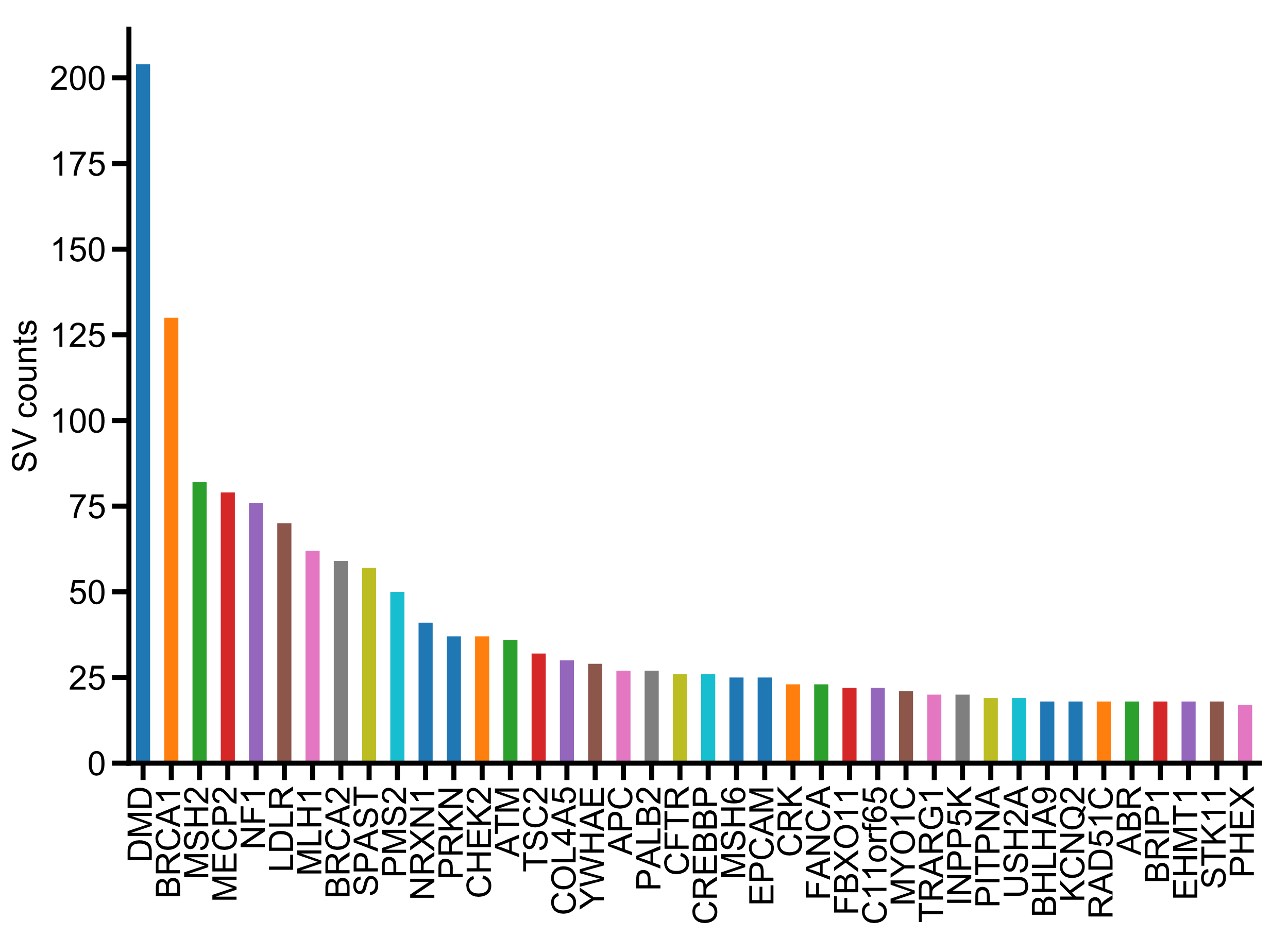

**Fig. S1.** Certain well-studied genes were overrepresented in our dataset. This plot shows the number of SVs overlapping each gene in the pathogenic ClinVar SVs (only showing the top 40 genes). For context, the median number of SVs overlapping each gene in our training dataset of pathogenic ClinVar SVs is 1.

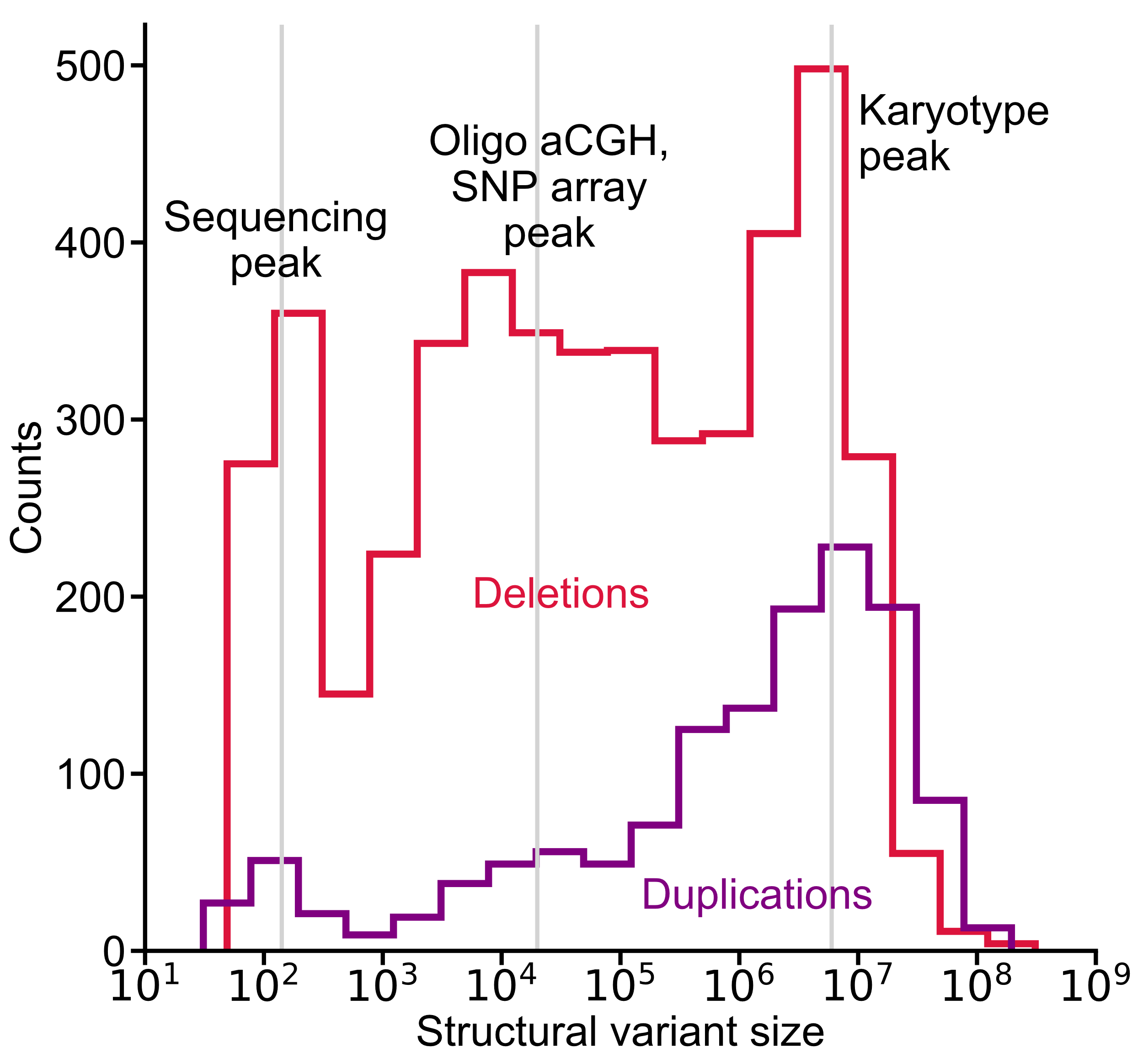

**Fig. S2.** Histogram of pathogenic SVs shows acquisition bias in SV size. Pathogenic SVs collected from ClinVar display three peaks consistent across both deletions and duplications. These peaks roughly correspond to the sensitivity range for three major methods to identify SVs: sequencing, oligo aCGH/SNP array, and karyotyping.

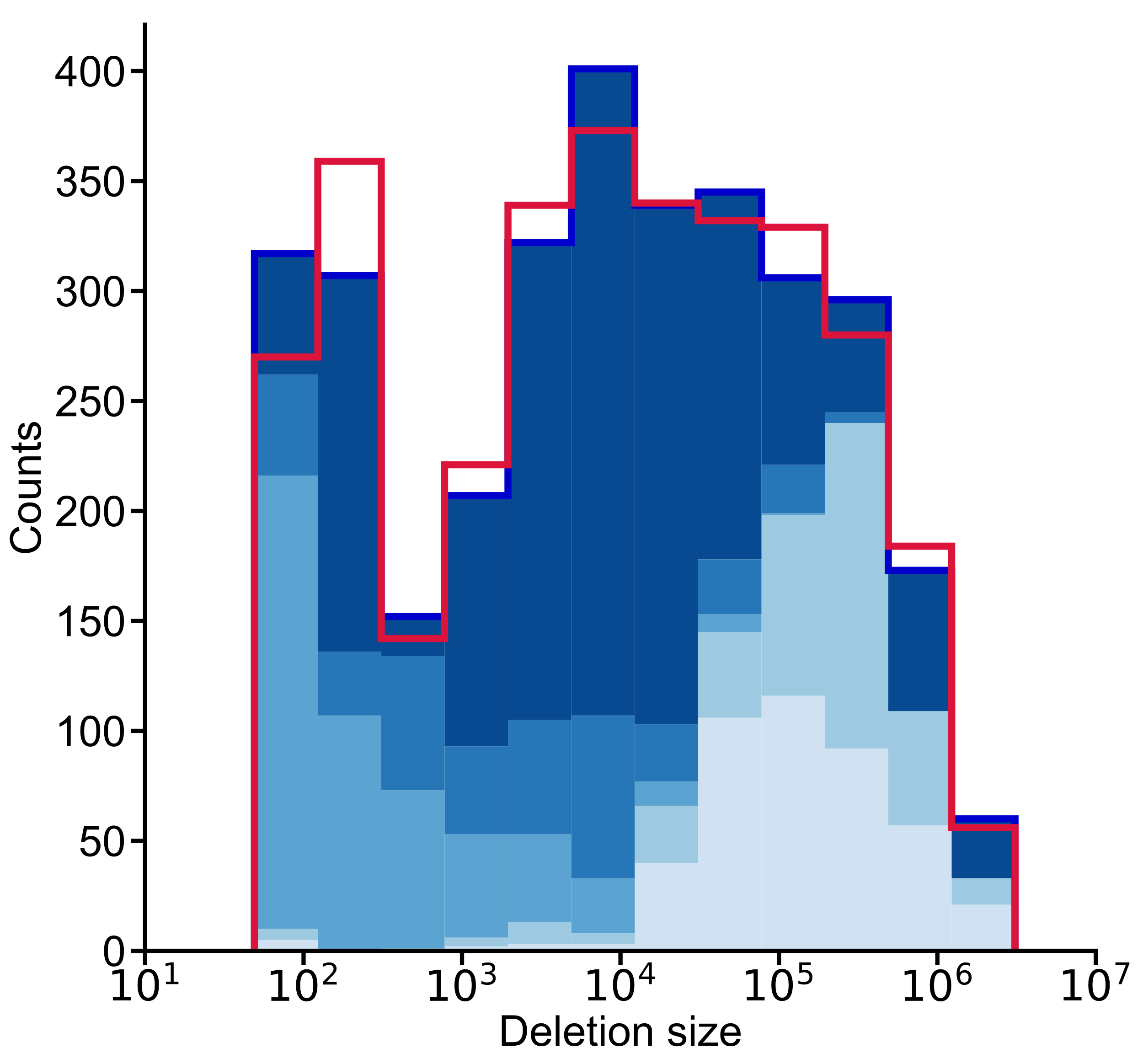

**b**

**a**

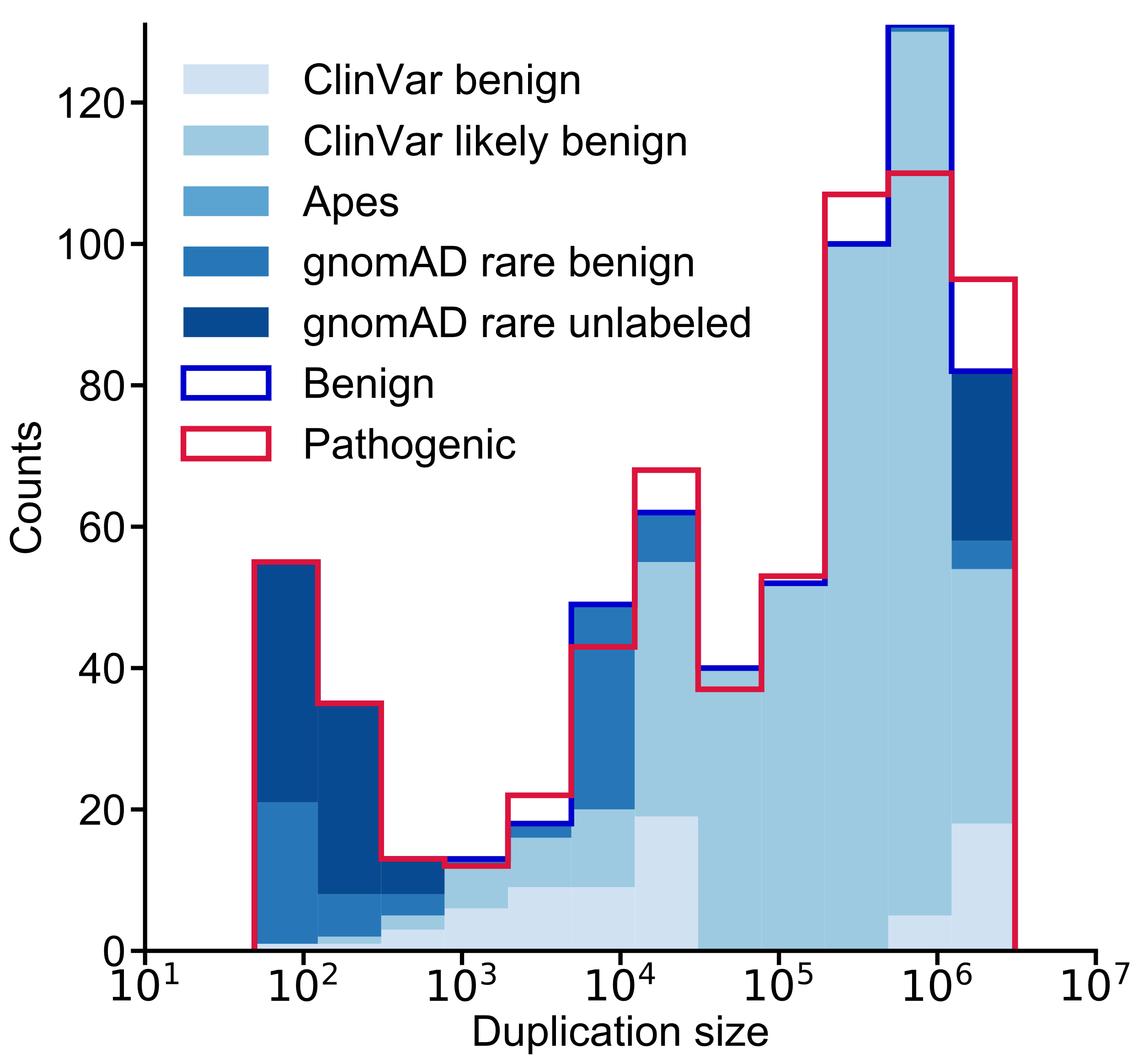

**Fig. S3.** Drawing SVs from multiple datasets allowed StrVCTVRE to train on a balanced dataset of SVs across a large size range. This is a more detailed version of Fig. 3. **a** Benign deletions were composed of 14% ClinVar benign, 12% ClinVar likely benign, 16% apes, 12% gnomAD rare benign, and 46% gnomAD rare unlabeled. **b** Benign duplications were composed of 11% ClinVar benign, 64% ClinVar likely benign, 11% gnomAD rare benign, and 14% gnomAD rare unlabeled.

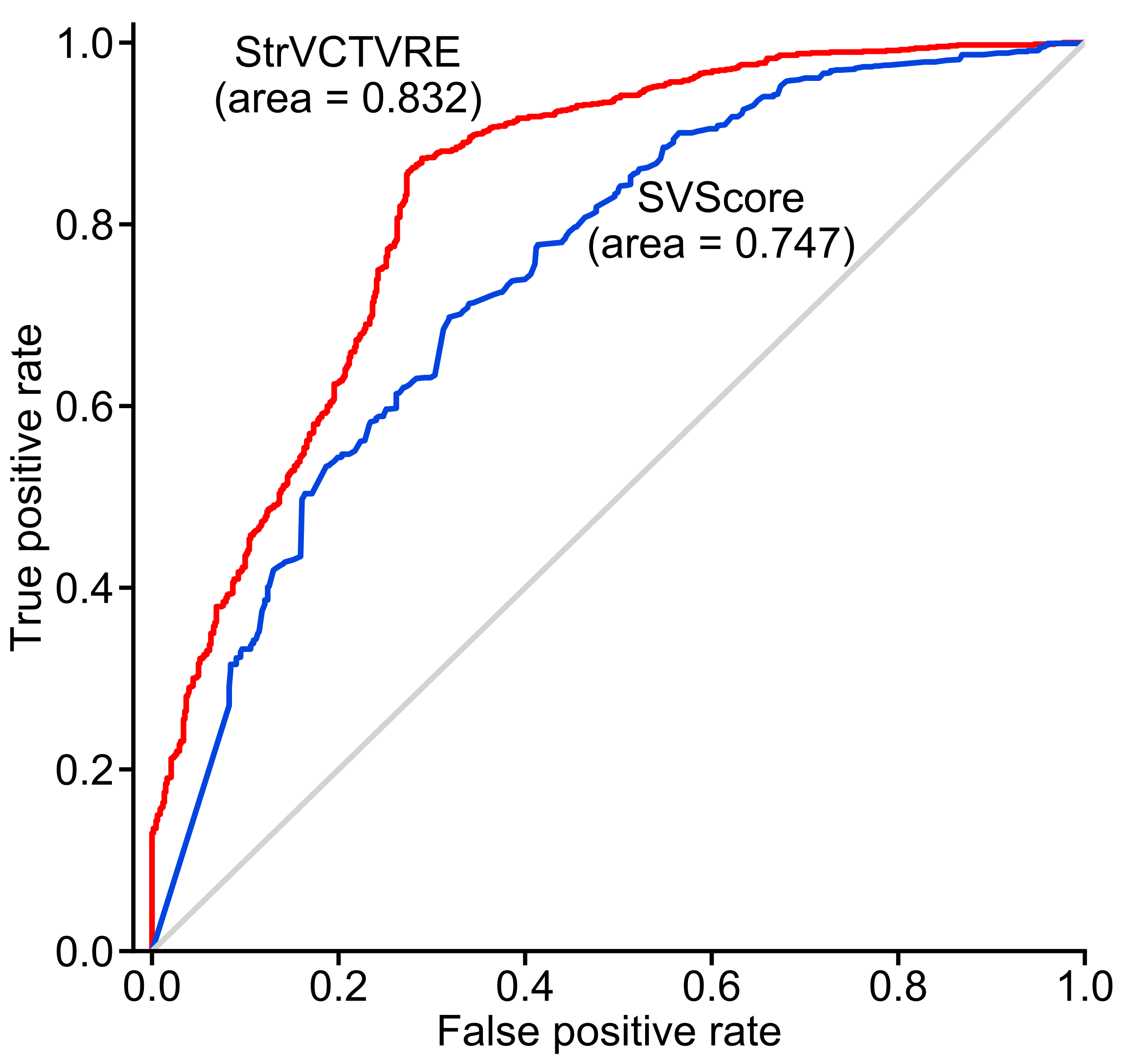

**a**

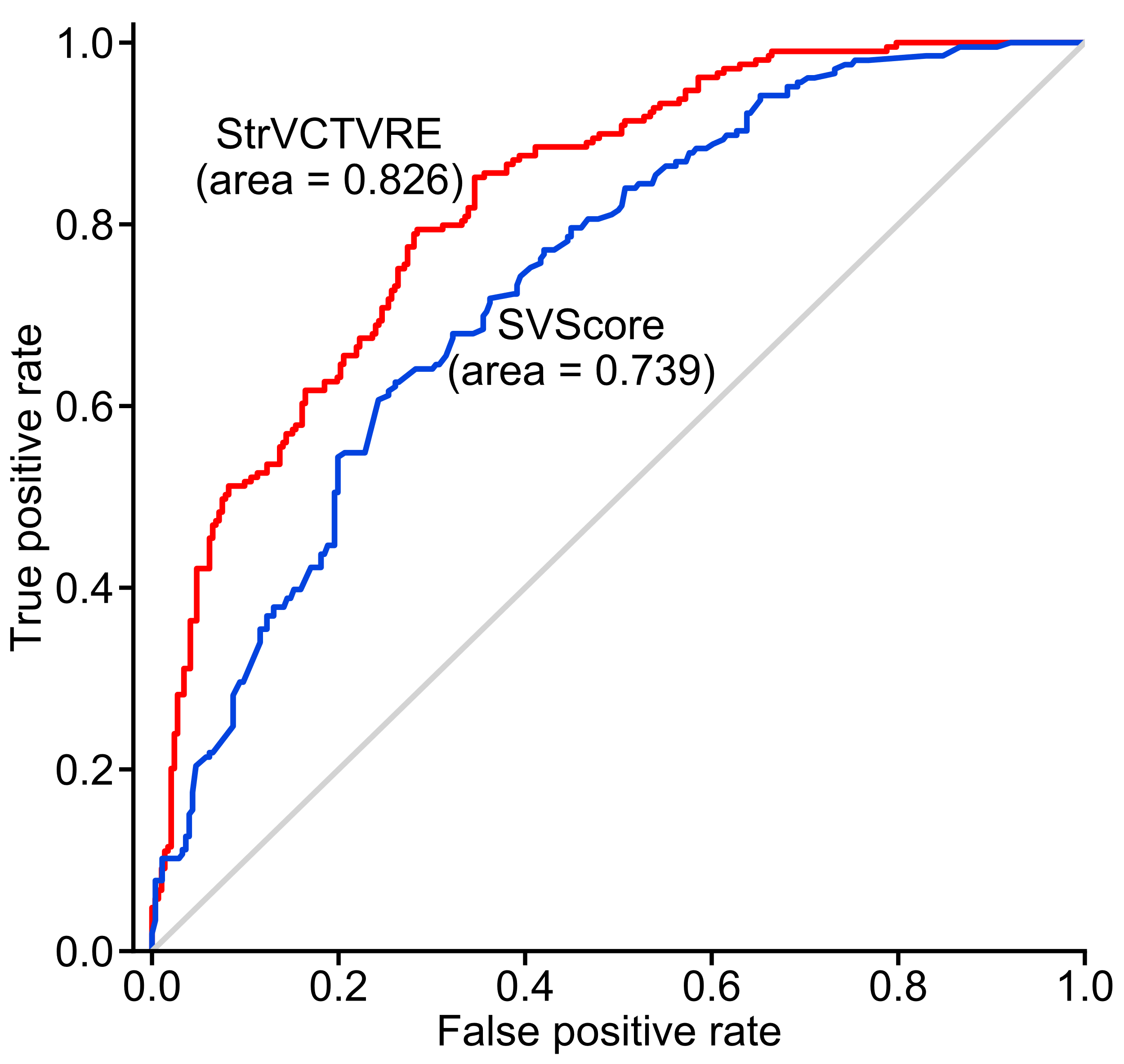

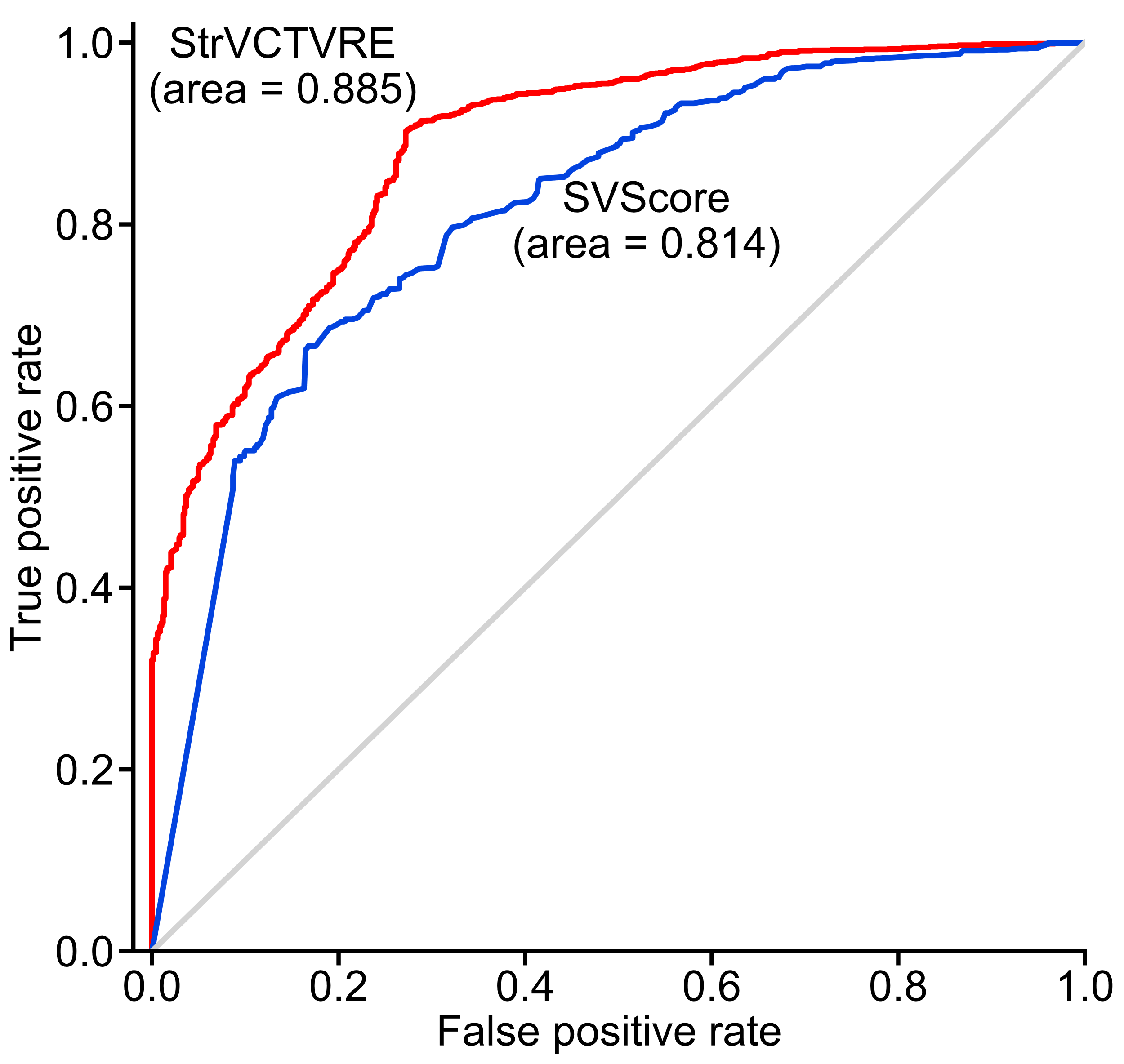

**c**

**b**

**Fig. S4.** StrVCTVRE and SVScore performance comparison on various held-out test datasets (ClinVar SVs on chromosomes 1, 3, 5, 7). **a** Duplicates and common variants were not removed from the test data. **b** Duplicates and common variants were not removed from the test data, and no size limit was imposed. **c** Duplicates and common variants were removed from the test data, and only SVs smaller than 1 Mb were included.

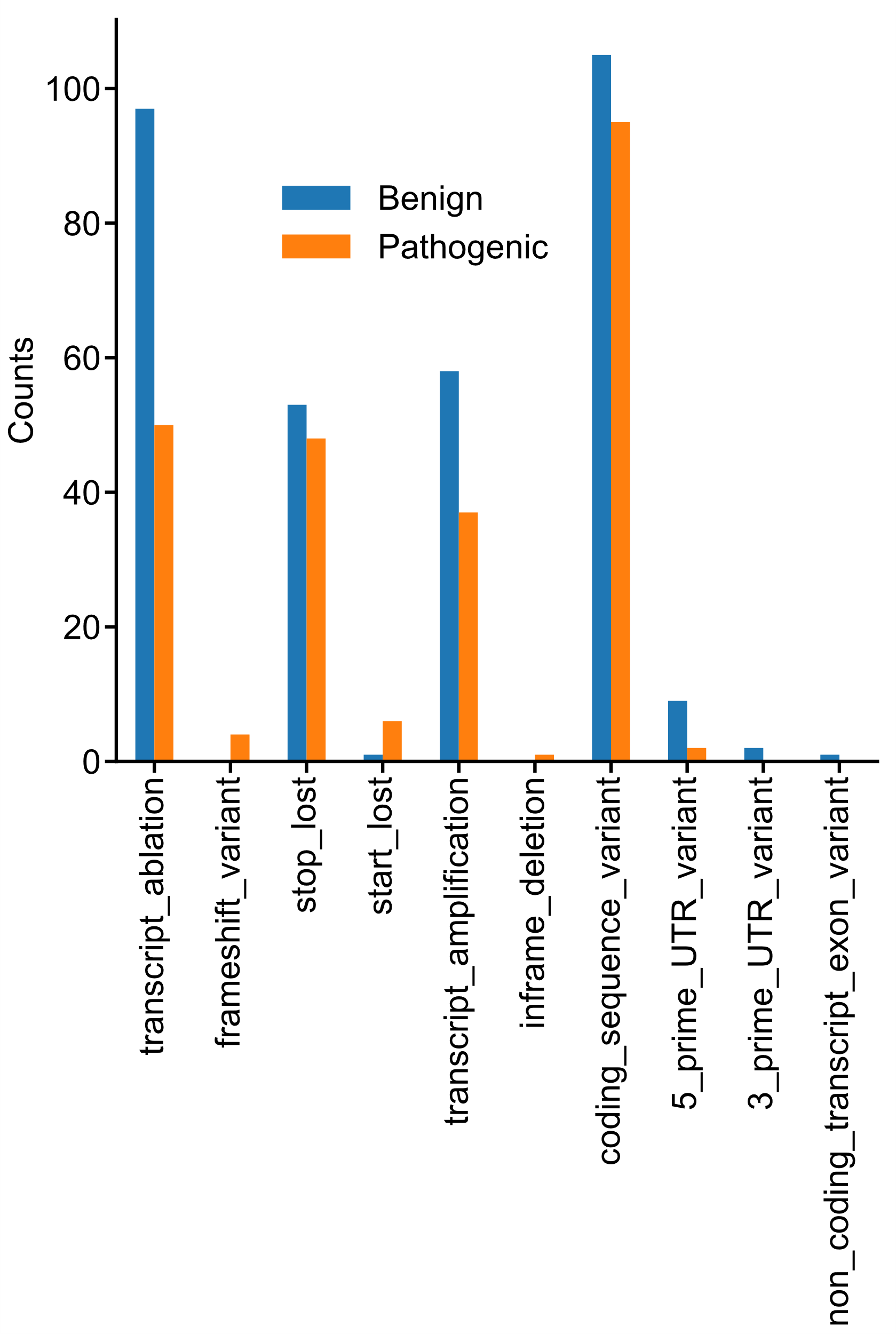

**Fig. S5.** Number of ClinVar test SVs (chromosomes 1, 3, 5, 7; from Fig. 2b) that VEP annotated with various sequence ontology terms. VEP classified more benign SVs than pathogenic SVs as transcript_ablation, leading to its poor performance in SV classification. Categories are ordered from left to right in descending deleteriousness.

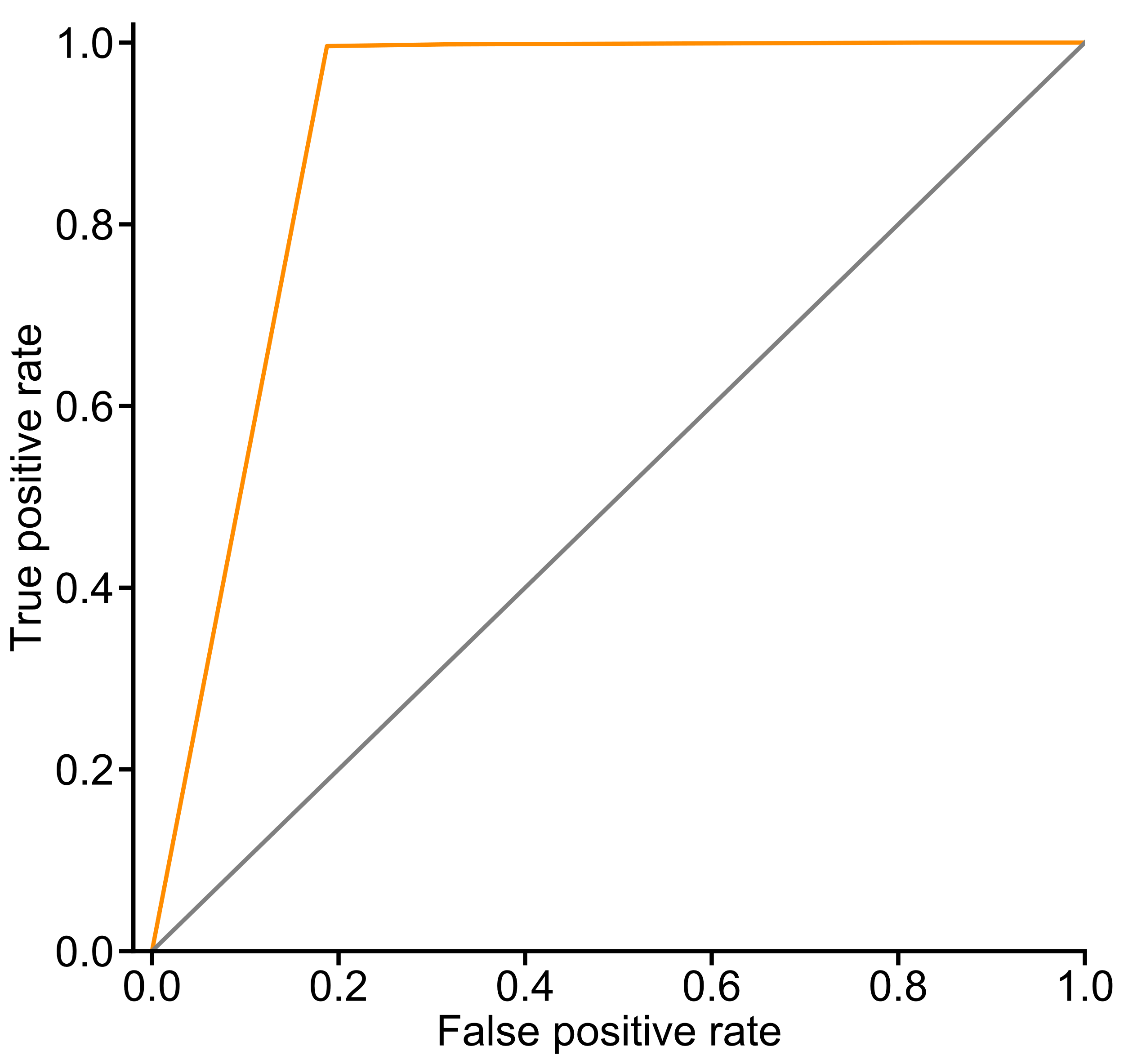

**Fig. S6.** AnnotSV excelled when tested on ClinVar variants that have a high degree of overlap with its cataloged variants. AnnotSV (orange) has an AUC of 0.905 when tested on 1042 ClinVar pathogenic variants and 867 ClinVar benign variants. It predicts nearly all pathogenic variants as pathogenic, along with 19% of the benign variants. The gray line represents chance.

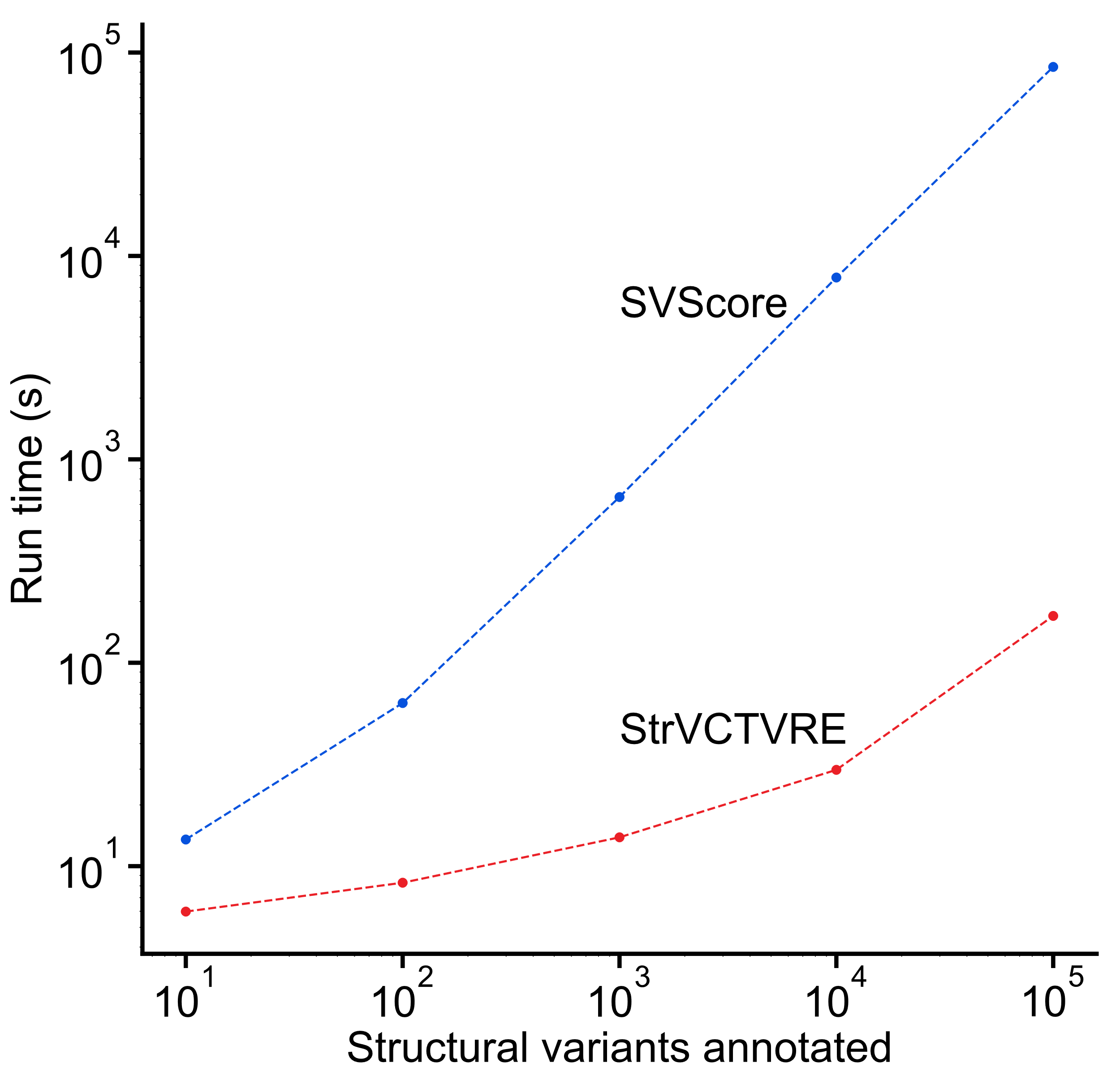

**Fig. S7.** Comparison of time required for StrVCTVRE and SVScore to annotate the same N structural variants. Each method was run on a 64-core Linux server with 2.6GHz Intel Xeon CPUs.

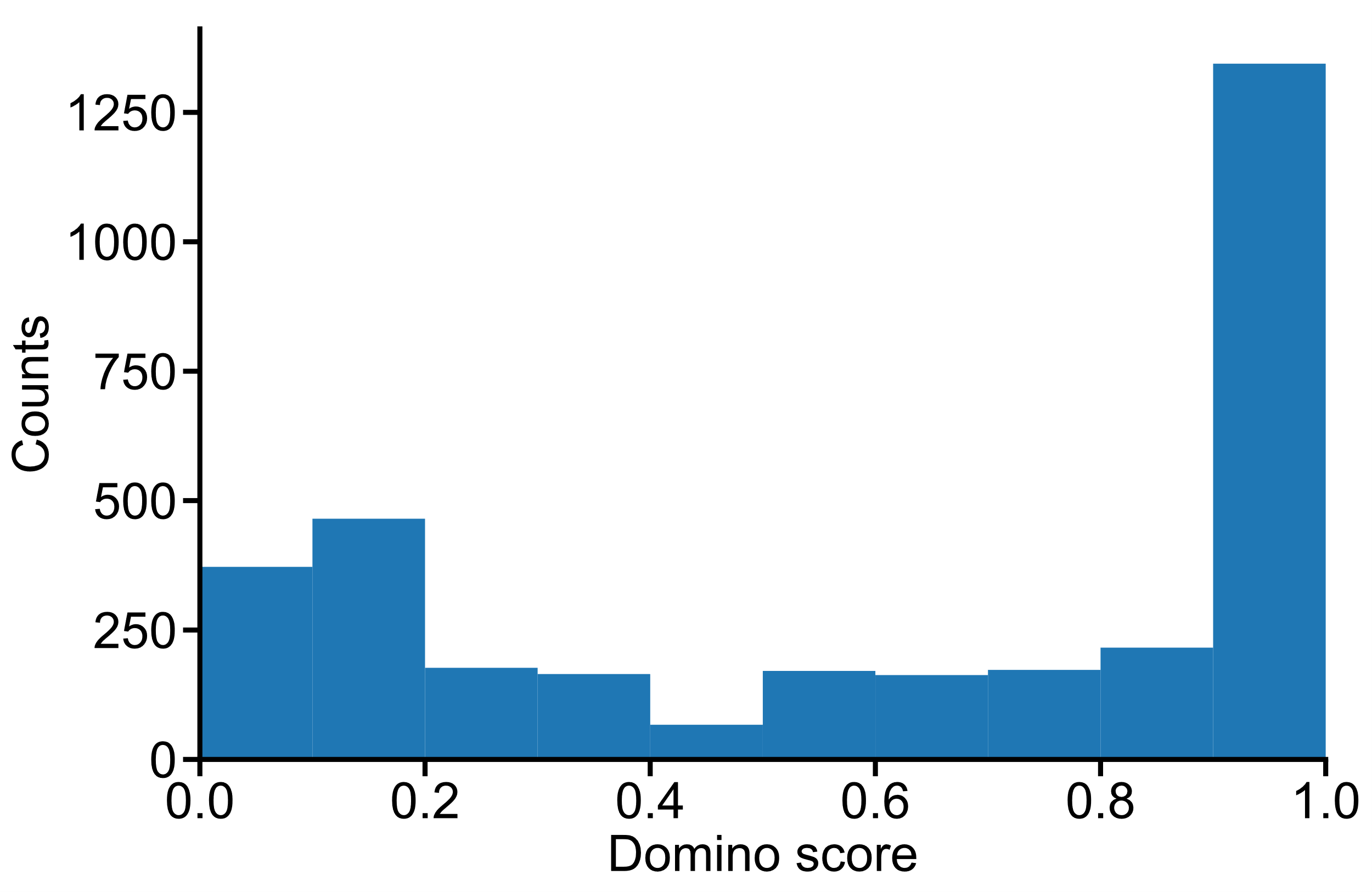

**Fig. S8.** Histogram of the predicted dominance of pathogenic SVs used in training. Predicted dominance was calculated for each SV by taking the maximum Domino(1) score of all genes the SV overlaps. SVs with larger Domino scores are more likely to have a dominant effect. Using a threshold of 0.5, 62% of pathogenic SVs are predicted to have a dominant effect. This analysis excludes SVs on sex chromosomes as they are not scored by Domino.

**b**

**a**

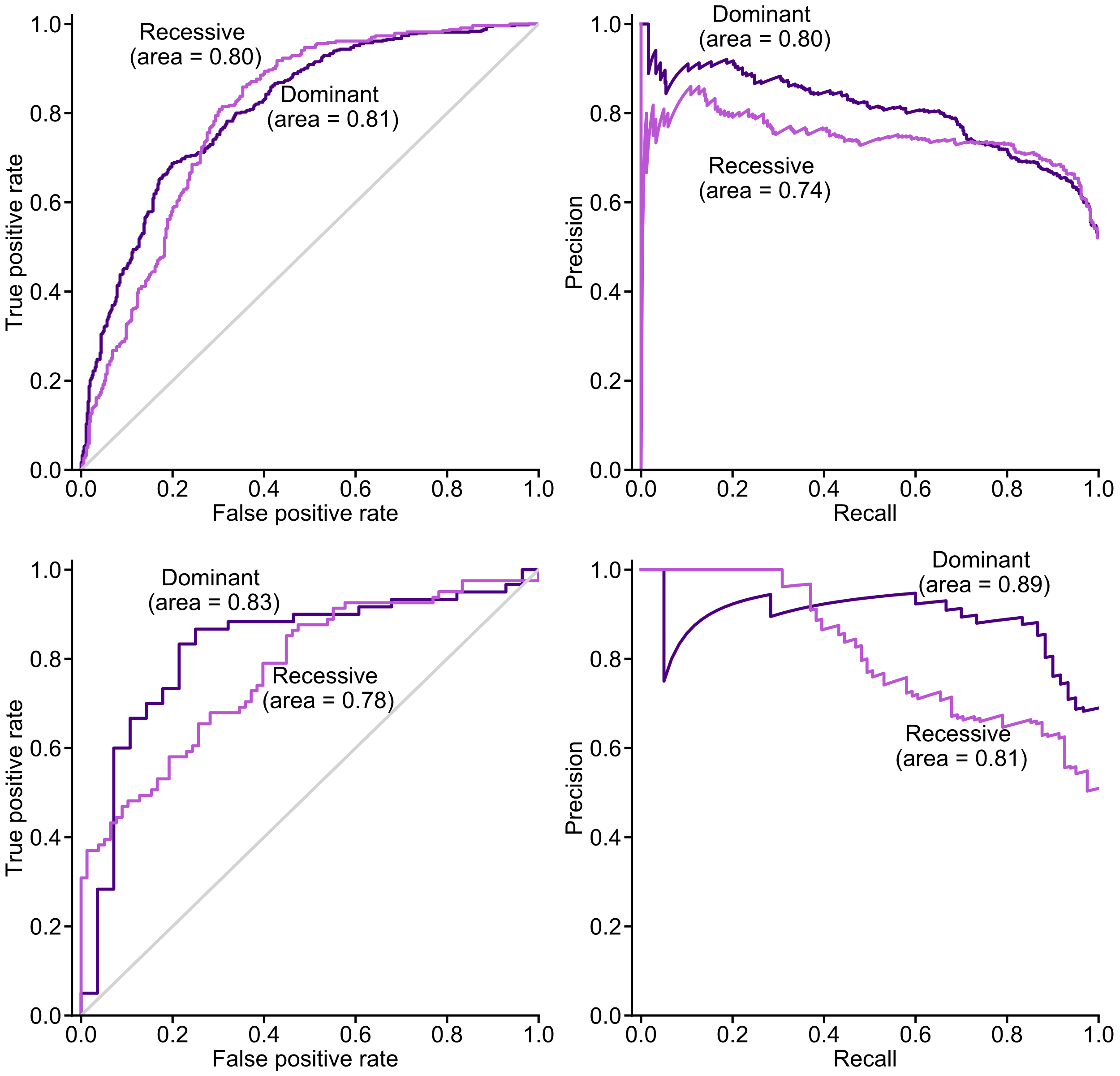

**d**

**c**

**Fig. S9.** ROC and Precision-Recall Curve (PRC) plots comparing StrVCTVRE’s performance on recessive and dominant SVs. **a** ROC and **b** PRC plot of StrVCTVRE’s performance on SVs in predicted dominant and recessive genes, using a Domino(1) threshold of 0.5 to separate dominant from recessive genes. **c** ROC and **d** PRC plot of StrVCTVRE’s performance on SVs overlapping high-confidence recessive or dominant genes identified by Balick et al(2). Overall, StrVCTVRE performs similarly on SVs in both dominant and recessive genes. There may be differences in performance at low sensitivity/recall but they are not consistent across datasets.

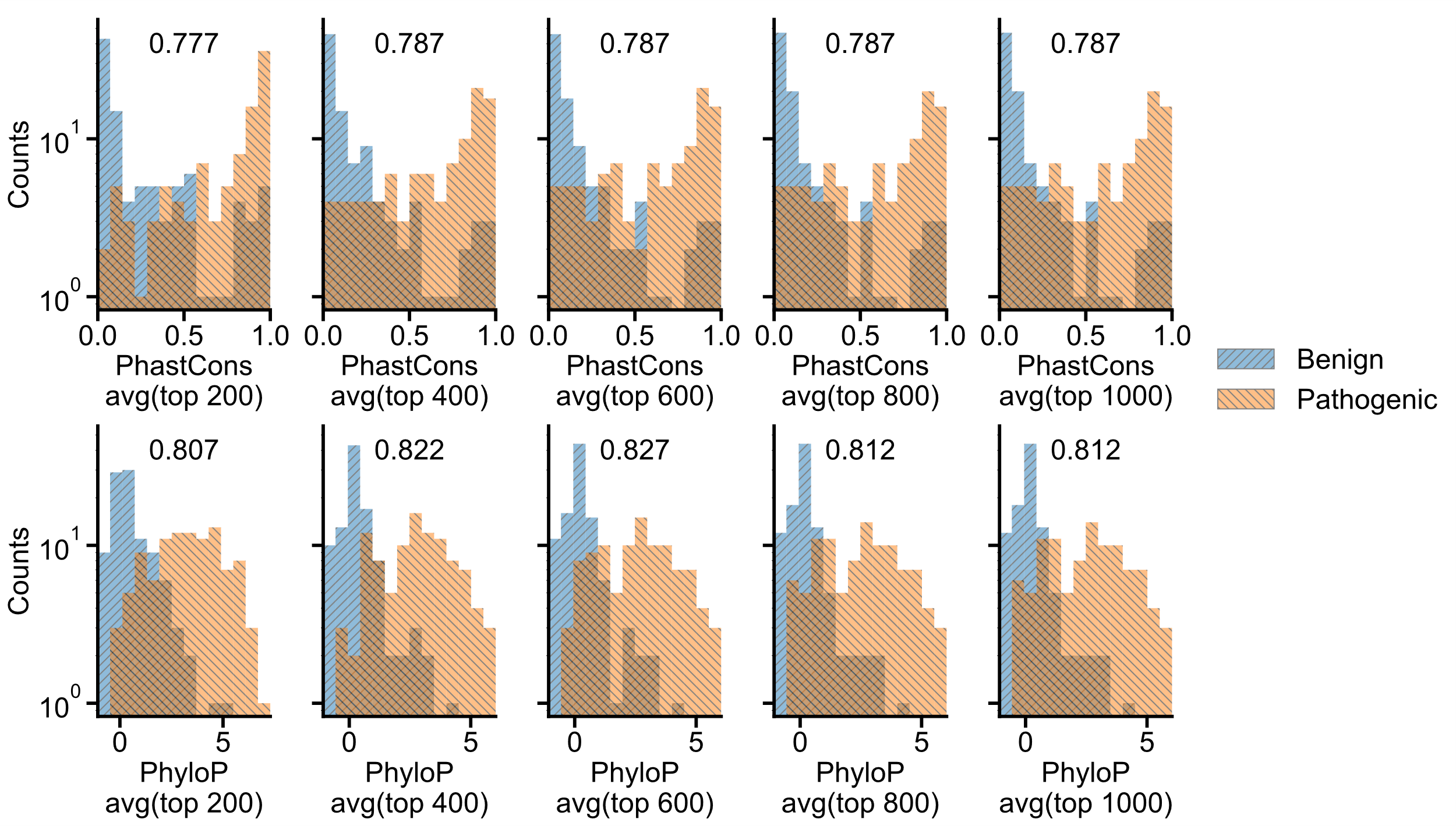

**Fig. S10.** Average of the top 400 PhyloP scores is one of the best indicators of SV pathogenicity. Comparison of PhyloP and PhastCons in differentiating between pathogenic and benign SVs. The columns represent different values of N, and the top row is PhastCons, while the bottom row is PhyloP. In each plot, the classification accuracy of a linear classifier is shown, and was calculated using a single feature: the average of the largest N PhyloP or PhastCons values among all positions overlapped by the SV. Each plot uses the same underlying SVs and shows a histogram of SV features.
